# Protection against disseminated disease following reinfection with *Mycobacterium tuberculosis* requires CD8+ lymphocytes

**DOI:** 10.64898/2026.10.29.762796

**Authors:** Andrew W Simonson, Michael C Chao, E Grace Bothwell, Feng Shan, Mark A Rodgers, Luke E Hood, Forrest Hopkins, Michael R Chase, Alanna Callendrello, Alexander A Smith, Ryan J Kelly, Cassaundra L Ameel, Edwin Klein, Jennifer Schober, Kara Kracinovsky, Jaime A Tomko, Harris Chishti, Marshall Malin, H Jacob Borish, Pauline Maiello, Charles A Scanga, Philana Ling Lin, Alex K Shalek, Sarah M Fortune, JoAnne L Flynn

## Abstract

Concomitant immunity is protective adaptive immunity conferred by pre-existing infection against subsequent exposures. This phenomenon is effective in the context of human tuberculosis (TB) and has been successfully recapitulated in non-human primates (NHPs). We previously demonstrated that depletion of CD4 T cells prior to secondary challenge with *Mycobacterium tuberculosis* (Mtb) only partially abrogated protection, implicating a critical role for additional cell types. Here, we describe the effects of depleting CD8α expressing cells in this model. CD4+ T cells in the airway were unable to compensate for lost cytotoxicity due to CD8 depletion. Contrary to the results of CD4 depletion, CD8α depleted animals exhibited a unique inability to limit Mtb dissemination, with significant lymph node involvement despite a short 4-week infection. These data suggest that concomitant immunity relies on non-redundant roles played by CD8+ lymphocytes. Thus, future vaccines should be designed to induce effective responses by both CD4+ and CD8+ populations to maximize efficacy.

**ONE SENTENCE SUMMARY:** Following reinfection with *Mycobacterium tuberculosis*, disease progression and dissemination is well controlled, however CD8+ lymphocytes play a unique and non-redundant role in containing spread to lymph nodes.

## INTRODUCTION

In humans, concomitant immunity provides effective protection against active tuberculosis (TB) following re-exposure to *Mycobacterium tuberculosis* (Mtb).^1^ Challenges with determining exact exposure timeline, dose and frequency, as well as distinguishing multiple infections, limit precise quantification of the impact on preventing TB disease in humans.^2^ The estimated 80% protection provided by prior Mtb infection outperforms the only vaccine currently in use, so the immune responses generated by initial natural infection may offer valuable insights for future vaccine design. However, the underlying immune mechanisms conferring this protection remain unclear in the context of TB.

A robust model of concomitant immunity has been established in cynomolgus macaques where Mtb infection timing and dose can be controlled.^3–5^ Serial PET CT imaging and genetically barcoded Mtb strains allow infection outcomes to be attributed to distinct challenge events.^3,5–7^ Animals with pre-existing Mtb infection developed significantly fewer granulomas following subsequent challenge with the same pathogen, with sterilizing immunity achieved in some cases.^3^ A subsequent study indicated that clearance of the primary infection with antibiotics partially abrogates the protective effects of prior infection.^4^ This protection is robust, extending across cases of reinfection with heterologous strains.^5^

We have sought to elucidate the immune mechanisms contributing to concomitant immunity against TB by interrupting the adaptive response generated by the primary infection. Previously, CD4 T cells were systemically depleted via antibody infusion prior to the secondary infection.^8^ This led to a partial loss of protection, with significantly increased bacterial burden in depleted animals compared to non-depleted reinfected animals, although disease did not progress to the extent seen in naïve controls. The impact of CD4+ T cell depletion on TB disease progression was primarily within the lung, with higher average bacterial burden in granulomas indicating the coordination of a competent immune response is impaired without these cells. Of note, the role of CD4 T cells in protection against reinfection appeared to be one of immune regulation and suppression of inflammation.^8,9^

Historically, in the field of TB immunology, the role of CD4 T cells, specifically Th1 cells producing IFN-γ, has been a primary focus.^10^ Relatively recently, more emphasis has been placed on how other immune cell types may be contributing to protection.^11–13^ The most obvious counterpart to CD4+ T cells is CD8+ lymphocytes.^14^ While conventional adaptive CD8+ T cells make up a sizable portion of this population, other cell types overlap functionally.^15^ This includes innate-like subsets, such as natural killer (NK) cells and cell types considered to bridge the gap between innate and adaptive responses, such as γδ T cells and donor unrestricted T (DURT) cells.

Here, we have targeted the CD8+ immune cells to determine whether all lymphocytes contribute redundant roles and functions, or if CD4+ and CD8+ populations are each uniquely necessary to prevent active disease following re-exposure to Mtb. Beyond recapitulating disease outcomes, NHPs have an immune system that more faithfully maps to that of humans.^16^ We employed antibody mediated depletion of all CD8α-expressing cells.^17,18^ Several cell populations are affected by this intervention, including the aforementioned lymphocyte subsets, but also a small proportion of dendritic cells.^15^ We demonstrated previously that depletion of CD8+ cells led to worsened outcomes in primary Mtb infection.^18^ Our results in the reinfection model indicate that CD8+ lymphocytes are essential to containing disseminated disease following secondary Mtb challenge, specifically with regard to lymph node involvement. Together with previous results in the same model indicating CD4 driven control of lung disease, these results highlight that although there are likely some functions shared between CD4+ and CD8+ lymphocytes, there are clearly non-redundant roles for these cell types in protection against Mtb reinfection.

## RESULTS

### Study design

Eighteen Chinese cynomolgus macaques (CCM) were intrabronchially infected with a low dose of a genetically barcoded library (Primary Library, or LibP) of Mtb (Erdman strain; median: 20.5 colony forming units (CFU)) (**Fig. 1A, Table S1**).^6^ This primary infection was allowed to establish for ∼ 9 weeks, affording ample time for mounting an adaptive memory response. Positron Emission Tomography – Computed Tomography (PET CT) scans were performed every four weeks to track progression, using fluorodeoxyglucose (^18^F) uptake as an indicator of inflammation which correlates with disease.^7^ Animals were then started on a standard regimen of rifampicin (R), isoniazid (H), ethambutol (E) and pyrazinamide (Z), or HRZE, for ∼ 18 weeks to clear the primary infection. While this is known to reduce the level of protection against a subsequent exposure, previous studies from our group have shown drug-treated animals are protected against TB disease following reinfection compared to naïve controls.^4^ It is essential to treat the primary infection prior to depleting lymphocyte subsets, since depletion would likely result in uncontrolled progression of the primary infection, limiting careful analysis of the secondary infection.^19^

**Figure 1.**
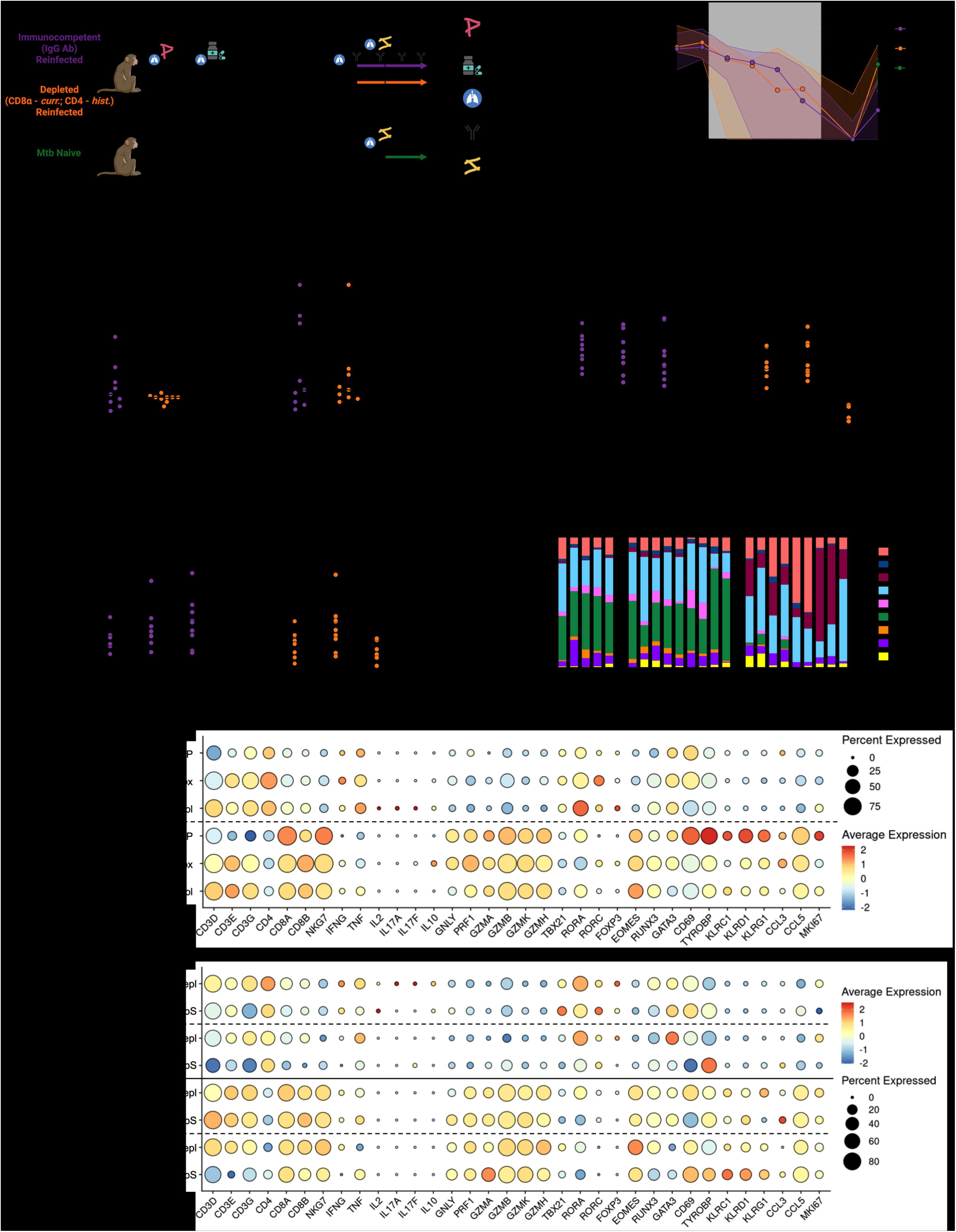
Study design and successful depletion in BAL. (**A**) Schematic showing the group designations, treatments, and sampling timeline. Made with BioRender. (**B**) Lung FDG activity quantified from serial PET CT scans during infections. Symbols represent group median; shaded regions represent 95% confidence interval. Grey region highlights period of drug treatment to clear primary (i.e. LibP) infection. (**C-D**) Number of granulomas identified by PET CT at (**C**) four and (**D**) eight weeks post-LibP challenge. P-values shown from Mann-Whitney non-parametric ranked sum test. (**E-H**) Antibody mediated depletion targeting CD8+ lymphocytes by BAL. Each symbol represents an animal; lines connect animals across timepoints. Frequency of CD8α+ T cells (**E,F**) shown as a proportion of CD3+ lymphocytes. Frequency of NK cells (**G,H**) shown as a proportion of CD3-CD20-lymphocytes. Animals received (**E,G**) an IgG isotype control antibody infusion or (**F,H**) an anti-CD8α depleting antibody infusion. Each symbol represents an animal; lines connect animals across timepoints. For panels E-J, p-values shown from repeated measures one-way ANOVA with Dunnett’s multiple comparisons test. (**I**) Dot plot showing frequency and level of expression for common T cell genes before, during and after primary infection with Mtb. (**J**) Relative abundance of lymphocyte subsets prior to secondary (i.e. LibS) infection. Each bar represents a BAL from one animal. (**K**) Dot plot showing frequency and level of expression for common T cell genes before and after infusion with control (IgG) or depletion (α-CD8α) antibodies.

The macaques were randomized into two groups (n = 9 each) based on total lung FDG activity via PET, a proxy for inflammation due to Mtb infection, 8 weeks post-LibP infection: anti-CD8α depletion or non-depleted (receiving IgG isotype control antibody). Serial PET CT scans were performed monthly and any disease (e.g. granulomas, consolidations, pneumonia) was annotated for tracking over time. Total lung inflammation (FDG activity above background) was quantified from the scans (**Fig. 1B**). Lesions identified on scans prior to the LibS challenge were attributed to the primary infection, enabling differentiation from subsequent disease at necropsy. Following randomization, the IgG and anti-CD8α groups had similar median numbers of granulomas and total lung FDG prior to antibiotic treatment (**Fig. 1B-D**). Beginning 2 weeks prior to the secondary challenge, these groups were intravenously infused with the assigned antibody every two weeks until necropsy. Both groups were intrabronchially challenged with a second, distinct genetically barcoded library of Mtb (Erdman strain; Secondary Library, or LibS; median: 11 CFU) for four weeks before necropsy.^3,6,8^ A third group (n = 5) of naïve macaques (i.e. no prior exposure to Mtb) were infected with LibS in parallel.

### Airway lymphocytes are expanded and express cytotoxic functions following primary Mtb infection

Serial bronchoalveolar lavages (BAL) were performed throughout the primary infection phase (**Fig. 1A**). The IgG and anti-CD8α groups had similar levels of CD8+ T cells, CD4+ T cells and NK cells during primary infection (i.e. prior to depletion) (**Fig. 1E-H, Fig. S1A,B**). Although the relative abundance of airway lymphocytes did not seem to be altered by prior infection (**Fig. S1C,D**), lymphocyte populations present in the airway, including CD8αβ T cells, CD4 T cells and NK cells, significantly increased in number following primary infection (**Fig. S1E-L**).

Single-cell RNA sequencing (scRNA-seq) and flow cytometry were used in parallel to assess the impact of primary Mtb infection, drug treatment and CD8α depletion on airway T cells (**Table S2**). Using the Seq-Well S^3^ platform for scRNAseq,^8,18^ overall recovery was consistent across samples in each experimental group (**Fig. S2A**). All expected cell types were observed and represented across experimental conditions (**Fig. S2B,C**). Monocytes and macrophages dominated the airway microenvironment (**Fig. S2D**). A well-defined T/NK population was isolated via coarse level annotation followed by two rounds of sub-clustering to remove contamination signal, allowing for identification of CD4-like, CD8-like and NK-like subsets (**Fig. S2E-G**).

Within eight weeks of initial exposure to Mtb, CD4-like T cells showed more frequent expression of *GZMB*, with modest changes in *GNLY* and *PRF1* (**Fig. 1I**). The increase in *GZMB* expression by CD4-like T cells paled in comparison to even baseline levels for CD8-like cells. Within CD8-like T cells, we observed strong upregulation of *PRF1* following Mtb infection. This cytotoxic transcriptional profile was accompanied by increased *EOMES* and *RUNX3* expression. Similar functional changes were observed by flow cytometry in airway lymphocytes stimulated with Mtb whole cell lysate (WCL) following Mtb exposure, with significantly more cells exhibiting effector functions following primary infection (**Fig. S3**). The number of CD8αβ T cells, CD4 T cells, and NK cells producing cytotoxic effectors significantly increased (**Fig. S3G-I**). The number of CD4 and CD8αβ T cells in airways were relatively equivalent before (median CD4: 4.6 log_10_, CD8αβ: 4.6 log_10_) and after (median CD4: 5.0 log_10_, CD8αβ: 5.3 log_10_) primary infection. However, more CD8αβ T cells produced granzyme B (0.95 log_10_ higher) and K (0.6 log_10_ higher) than CD4 T cells at baseline (pre-primary infection). This widening functional gap post-primary infection was seen in both granzyme B (1.3 log_10_) and granzyme K (0.9 log_10_) production. In contrast, CD4 T cell functionality was highlighted by significant increases in IFNγ, TNF, IL-2, and IL-17 production. This was corroborated by increased *TNF* and, to a lesser extent, *IFNG* expression by CD4-like T cells. *RORA* and *RORC* upregulation observed in BAL post-Mtb infection supports flow cytometric data demonstrating enriched Th17 populations (**Fig. 1I**). RORα expression has also previously been detected by intranuclear staining and RNA sequencing of granuloma T cells.^20,21^ Significantly more CD8αβ T cells and NK cells produced IL-2 and IL-17 (**Fig. S3J-L**). IL-17 production by airway NK cells was also observed in IV BCG vaccination.^17^ Both transcriptomic and flow cytometric evidence supports functional specialization between CD4 T cells and traditional cytotoxic lymphocytes. Together, these data suggest strong recruitment of several types of lymphocytes to the airway following primary infection, with inflammatory functions poised to mount a rapid antimicrobial response. Functional remodeling of the immune environment in the airway was also a key feature found following IV BCG vaccination, presenting a common theme across successful protection models.^22^

Some of the functional shift initiated by primary Mtb exposure were lost following treatment with antibiotics (**Fig. 1I**). This is most likely related to reduced infection.

### CD8+ populations are successfully depleted in the airway

Antibody mediated depletion specifically targeted CD8α expressing cell types with expected declines in cell numbers (**Fig. 1E-H,J, Fig. S1C-K**). Depletion of CD8+ T cells was more effective than that of NK cells, an expected result since NK cells are defined by expression of at least one of the proteins CD8α, NKG2A or CD16, rather than the former alone (**Fig. 1E-H,J, Fig. S1**).^17^ As seen in previous depletion studies, following CD8α depletion, the relative abundance of CD8α-cell types (e.g. B cells, CD4+ T cells, innate lymphoid cells) expanded (**Fig. S1A-B**).^18^

Transcriptomic analysis of T cells prior to and after antibody mediated depletion indicated that the randomized IgG and α-CD8α groups had similar immune profiles in the airways following treatment of the primary infection (**Fig. 1K**). CD8-like T cells did not show substantial transcriptional changes following depletion, indicating that all subsets (e.g. functional and non-functional) were depleted evenly. No notable changes were seen following administration of the IgG isotype control antibody. Following infusion with α-CD8α depletion antibody, the CD4-like T cells adopted a compensatory cytotoxic profile, with more frequent and abundant *GZMB* expression and downregulation of *IFNG* and *TNF*. We observed the same for the corresponding transcription factors, with reduced expression of *TBX21*, *RORA* and *GATA3* and increased *RUNX3* expression. These data align with previous scRNA-seq findings from a six week primary infection with Mtb following CD8α depletion in which CD4 T cells compensated for some (e.g. *GZMB*), but not all (e.g. *PRF1*), cytotoxic function missing in the absence of CD8 T cells.^18^

### CD8α depletion partially abrogates protection against TB pathology

To improve interpretation of the current study, we compare these results to data adapted from our previous CD4 depletion study in our reinfection model on the right side of relevant graphs.^8^ Data previously reported from that study are shown on a grey background, while new analyses of data collected during that study do not have a background added. No statistical tests were performed by comparing across studies; the plots are intended only for qualitative comparison and context.

FDG PET signal can be quantified as a surrogate for disease.^7,23^ At necropsy, CD8α depleted animals had similar levels of lung inflammation assessed by PET to the naïve group and had significantly more inflammation compared to IgG control animals (**Fig. 2A**). Serial PET CT imaging throughout both infections enabled tracking of lesions identified prior to or following LibS challenge (**Fig. 2B**). Scan-based labeling was then validated via Mtb barcode sequencing to establish which library or libraries are present in each sample. Due to drug treatment of primary infection in this study, no LibP barcodes were found in any samples. Thus, all bacterial burden and tissues involved in active disease are attributed to the secondary challenge (**Table S3**). Despite a broader range of outcomes, similar to those seen in naïve animals, the number of granulomas attributable to the secondary (LibS) infection were not significantly different across the groups in this study (**Fig. 2C**). Unique LibS barcodes per animal, a measure of the number of inoculated bacteria able to establish infection, mirrors the slightly broadened range of numbers of new granulomas in CD8-depleted animals (**Fig. S4A**). These data suggest that both CD4 and CD8+ lymphocytes impose a small bottleneck on establishment of infection following re-exposure.

**Figure 2.**
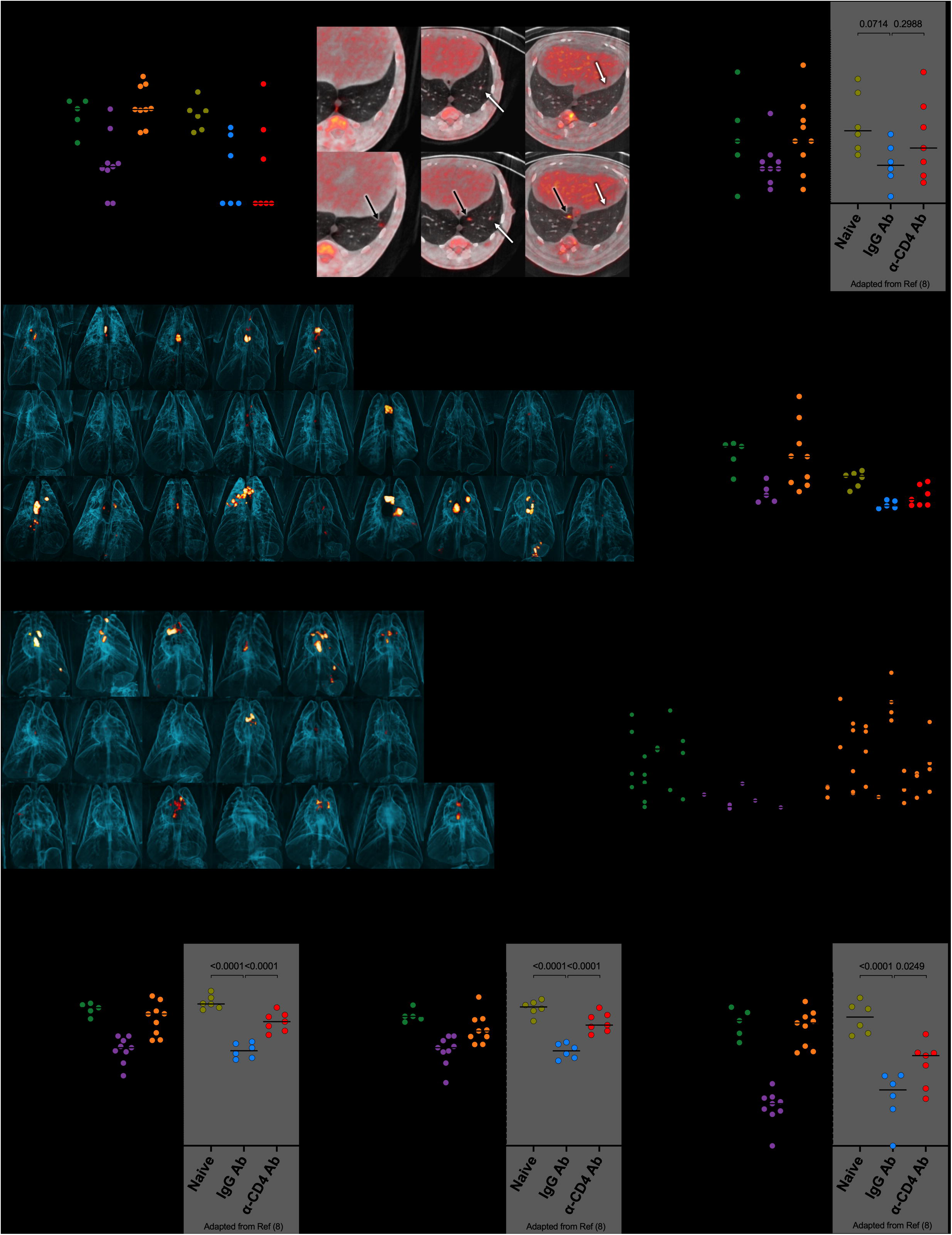
CD8α depletion leads to exacerbated TB disease driven by lymph node involvement. Data from the current study are shown alongside data adapted from a previous study from our group in which CD4+ cells were depleted in the same reinfection model.^8^ No statistical comparisons are made across studies and rather serve as qualitative comparisons to contextualize results. (**A**) Quantification of FDG activity in lung from pre-necropsy PET CT scan. (**B**) Representative slices from PET CT scans demonstrating annotation of lung granulomas, temporal tracking with serial scans and differentiating new and old lesions. White arrows indicate “old” lesions from primary LibP infection. Black arrows indicate “new” lesions that developed after the secondary LibS infection. (**C**) Number of granulomas found at necropsy that were attributable to LibS based on PET CT scans and genetic barcoding analysis. (**D,E**) 3-dimensional volume rendering of PET CT scans from (**D**) current CD8 depletion study and (**E**) previous CD4 depletion study. (**F**) Mean SUVR per LN identified on the pre-necropsy scan. Each symbol represents an animal, line represents group median. (**G**) SUVR per individual LN identified on the pre-necropsy scan. Each column represents an animal, each symbol represents a granuloma, line represents animal mean. (**H-J**) Bacterial burden attributable to LibS in (**H**) all thoracic tissues, (**I**) only lungs, including random samples from each lobe and isolated lesions and diseased tissue, or (**J**) thoracic lymph nodes. Statistics for data from the previous CD4 depletion study are copied from the published study if previously reported (grey boxes) or following the same methods as the current study for metrics not previously reported (no grey box). For panels A,F,H-J, p values shown are for Welch’s ANOVA with Dunnett’s T3 multiple comparisons test. For panel C, p values shown for Kruskal-Wallis non-parametric test with multiple comparisons. In CD8α depletion study, all groups were compared against each other. In CD4 study, all groups were compared to the non-depleted IgG reinfection control. Each symbol represents an animal; line represents group median.

### Functional NK cells in the airway correlate with early containment of Mtb infection

To elucidate which CD8a+ cell types might be associated with this modest early control phenotype defined by low establishment of barcodes in those with prior Mtb infection, we considered the size of innate-like CD8+ populations present in the airway. The CD8 depletion group provided a sample set with both a range of depletion levels and number of unique barcodes establishing infection. The IgG group, while more limited in the ranges of numbers of cell types and established barcodes, contextualizes these findings to the model of Mtb reinfection in immunocompetent individuals. The naïve group was too small to perform correlation analysis on but is plotted to highlight the differences in immune responses driving protection, or lack thereof. Previous work from our group identified the number of NK cells in the airway (via BAL) as a correlate of protection conferred by intravenous administration of BCG, another effective model of protection against Mtb infection.^24^ Here, NK cells, but not γδ T cells, CD8αα T cells, or CD8αβ T cells, were significantly negatively correlated with the number of unique barcodes recovered (**Fig. S4B-E**). While this correlation was not statistically significant in non-depleted animals, the data sets align well in overlapping regions, suggesting that the relationship may hold true across concomitant immunity models given larger sample sizes. This appears to be driven by pro-inflammatory functions, with IFN-γ and TNF producing NK cells being correlated with fewer established barcodes, and cytotoxic (i.e. granzyme producing) NK cells trending similarly (**Fig. S4F-K**). Notably, naïve animals (green) have consistently lower numbers of NK cells in the airways compared to previously infected animals and there is no relationship with establishment of barcodes (vertical distribution; **Fig. S4B**). However, the number of IFN-γ producing NK cells in BAL shows a strong relationship with barcode establishment across all three treatment groups, suggesting this is a critical population for early responses to Mtb (**Fig. S4H**).

### Dissemination to lymph nodes is exacerbated in absence of CD8+ lymphocytes

Substantial disease was observable on pre-necropsy PET CT scans in naïve and CD8α depleted animals (shown as three-dimensional reconstructions in **Fig. 2D**, **Fig. S5A**), with much less apparent disease in IgG controls. Similar PET CT scans in CD4 depleted animals appeared to show less disease (**Fig. 2E**).^8^ Individual granulomas are easily observed by PET CT by analyzing slices (**Fig. 2B**) but it can be difficult to distinguish them from uninvolved lung tissue in the 3D renderings. However, extensive disease manifestations, such as TB pneumonia, consolidations, and inflamed lymph nodes (LNs), appear as large PET hot structures (**Fig. 2D**,**E****, Fig. S5A**). A gross pathology scoring system, as published previously, quantifies evidence of disease found at necropsy, including remnants of previously treated (i.e. LibP) infection.^23^ Lung pathology scores were similar across the groups, in part due to evidence of prior Lib P pathology in the IgG and CD8-depleted group which increases the pathology score (**Fig. S5B**). In contrast, CD8α depleted animals had significantly more pathology in lymph nodes (evidence of granulomas and necrosis) than IgG controls, similar to LN pathology scores seen in the naïve group (**Fig. S5C**). The depleted group also had significantly higher levels of inflammation in thoracic LNs, as quantified by PET signal (**Fig. 2F**). In contrast, LNs in CD4 depleted animals remained similar to IgG control animals in terms of FDG+ thoracic LNs, implying a major difference in roles for CD8α+ and CD4+ lymphocytes in containing disseminated disease. Both the number of FDG-avid LNs and the level of inflammation (FDG uptake) within those tissues increased following CD8α depletion compared to IgG control animals (**Fig. 2G**).

At necropsy, all TB pathology (e.g. granulomas, other pathologies), random samples of uninvolved lung lobes, all thoracic lymph nodes, and peripheral lymph nodes were individually homogenized and plated for colony forming unit (CFU) enumeration. The total thoracic bacterial burden, quantified by summing CFU collected from all individual samples from lung (all granulomas, other pathologies, and uninvolved tissue from each lung lobe) and thoracic lymph nodes, was significantly higher (∼20-fold) in CD8α-depleted animals compared to IgG reinfection controls (**Fig. 2H**). A 14-fold increase was observed following CD4 depletion, and control groups (naïve and IgG) were at similar levels across studies. Depletion of CD4 and CD8α expressing cells led to an intermediate phenotype in the lung, with bacterial burden in those tissues indicating partial loss of protection (**Fig. 2I**). CD4 depletion led to a significant increase (∼20 fold over IgG, ∼50 fold below naïve) in LN bacterial burden attributable to the secondary challenge. In contrast, CD8α depletion resulted in >1000-fold increase in LN CFU compared to IgG controls; this was at the level of naïve animals (**Fig. 2J**). These data support PET CT and gross pathology data indicating significant loss of control of disseminated TB disease in lymph nodes in the absence of CD8+ lymphocytes. Concomitant immunity leads to a significant reduction in LN bacterial burden and the number of CFU+ (i.e. involved) LNs (IgG vs naïve, **Fig. 2J**, **Fig. 3A**). CD8α depletion eliminated protection at the level of LNs, while CD4 depletion had a minimal effect (**Fig. 3A**). The high lymph node burden was observed across multiple LNs in depleted animals, confirming this outcome does not stem from a single site becoming overwhelmed with disease (**Fig. 3B**).

**Figure 3.**
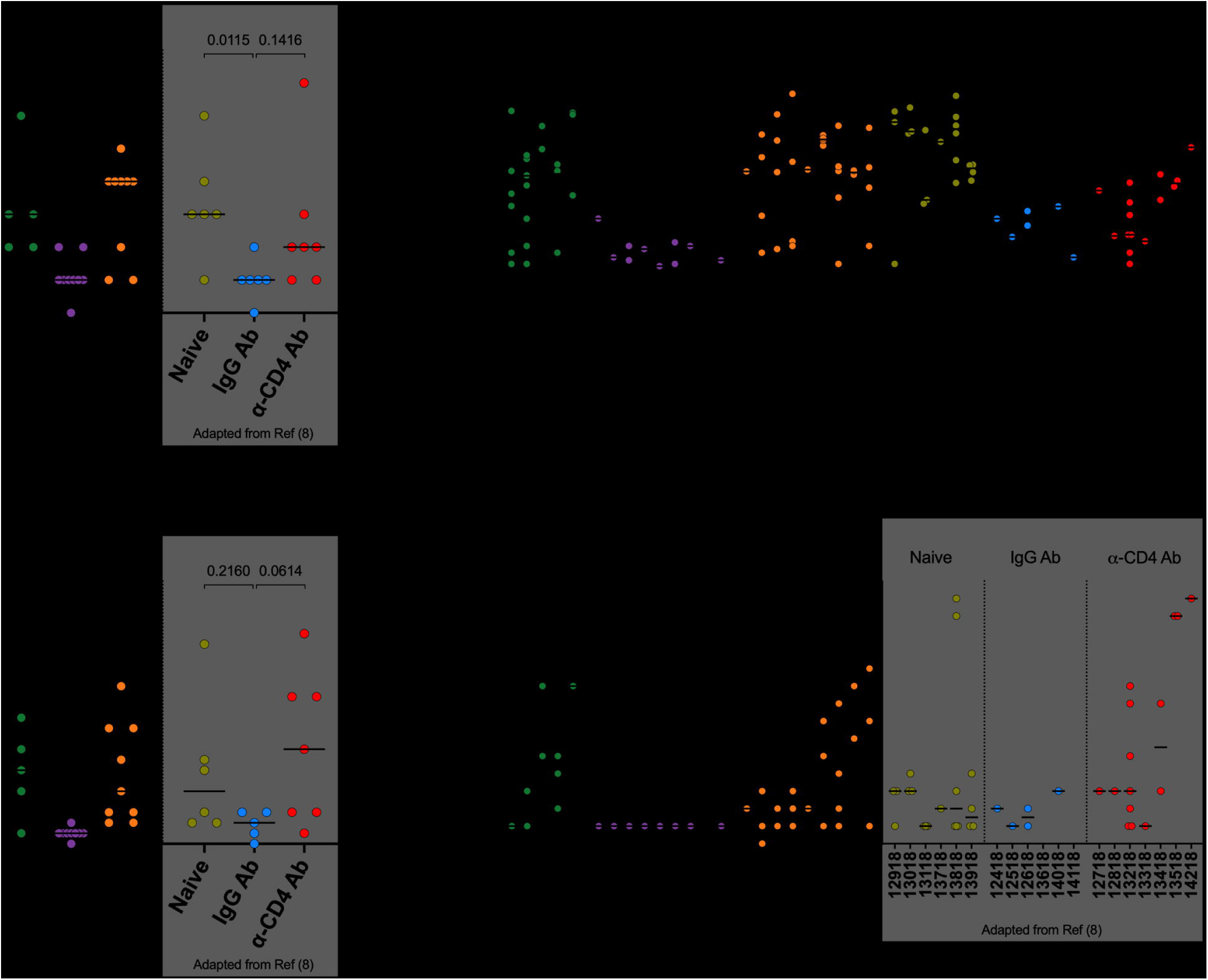
Analysis of dissemination to lymph nodes. (**A**) Number of involved lymph nodes that grew LibS Mtb. (**B**) LibS burden in each lymph node. (**C**) The mean number of unique barcodes recovered from CFU+ lymph node tissues. (**D**) Number of unique barcodes recovered from each lymph node. For panels A,C, each symbol represents an animal; line represents group median. For panels B,D, each column represents an animal, each symbol represents a lymph node; line represents animal mean. For panels A and C, p values shown are for Kruskal-Wallis non-parametric test with Dunn’s tests adjusted for multiple comparisons. In CD8α depletion study, all groups were compared against each other. In CD4 study, all groups were compared to the non-depleted IgG reinfection control.

The number of unique LibS barcodes identified per LN is another measure of dissemination; presumably this reflects spread from lung to thoracic lymph node. This is a normal process in infection, since T cells are primed by Mtb or Mtb antigens in lymph nodes. Thus, in naïve animals there is a wide range in the number of barcodes/LN, which is greatly reduced in IgG animals, another indication of protection (**Fig. 3C**,**D**). Depletion of CD8+ cells led to a significant increase in average number of unique barcodes in LNs, similar to naïve animals (**Fig 3C**,**D**). Thus, not only were more LNs involved in CD8α depleted animals with a higher CFU per site, but more individual dissemination events spread bacteria to the LNs in those animals. There was also a trend towards more LN dissemination in the context of CD4 depletion, though smaller group sizes in the prior CD4 depletion study are likely a factor in this effect on barcodes not reaching statistical significance. However, other metrics of LN involvement, such as bacterial burden, number of involved LNs, and inflammation, are skewed towards being influenced more so by CD8+ subsets. Thus, while both CD4+ and CD8+ lymphocytes contribute to preventing dissemination out of the lungs, CD8+ cell types appear to have a stronger effect on dissemination with more LNs harboring Mtb and appear to be uniquely capable of limiting bacterial outgrowth and disease progression at those sites. Our previous work in acute Mtb challenge showed CD8α depletion resulted in significantly higher CFU in LNs (p value = 0.0144), and CD8β depletion also showed a trend of an increase (p value = 0.0694), despite an early endpoint at just six weeks post challenge.^18^ These data further support the role of CD8+ lymphocytes in limiting disseminated disease.

### LN architecture is disrupted by extensive disease following CD8α depletion

Lesions that form in LNs disrupt the native lymphocyte organization and stromal architecture.^25–27^ If LNs do not control infection, granulomas can enlarge with increased necrosis, coalesce, eventually resulting in effacement of the entire LN with the potential to eventually erode into the airway and spread infection throughout the lungs. Given the increase in involved LNs observed in depleted animals, we next sought to determine whether the structural organization of involved tissue is affected more in the context of CD8α depletion. Unlike in lung granulomas, LN lesions can lack organized regions (e.g. a lymphocyte cuff) surrounding central caseum.

An uninvolved LN of a non-depleted (IgG) animal contained abundant populations of T cells (both CD3+CD8α– and CD3+CD8α+) in the paracortex without clear disruption, and minimal GrzB+ signal (**Fig. S6A**). Conversely, an involved LN from a naïve animal had large, multifocal lesions developing, indicated by displacement of T cells and strong GzmB+ cores (**Fig. 4A**). While an involved LN from an IgG animal showed clear disruption to lymphocyte localization, there was extensive staining of CD8α, much of it overlapping with CD3 staining indicating large numbers of CD8 T cells. However, there were minimal GzmB+ cells compared to naïve tissue (**Fig. 4B**). In LN tissue, more generally, there was a low frequency of CD8α+ GzmB+ cells, indicated by a lack of overlap between green and red channels (**Fig. 4A,B**, bottom middle, top right). Staining of an involved LN from a depleted animal showed an expected absence of CD8α+ cells throughout the tissue with multifocal lesions developing and areas of dense GzmB+ staining (**Fig. 4C, Fig. S6B**). This indicates that granzyme production is likely related to bacterial burden in the diseased area, possibly produced by neutrophils.^28^ Spatially, the arrangement of CD3+ cells in depleted tissue appears highly unorganized with less consistent density.

**Figure 4.**
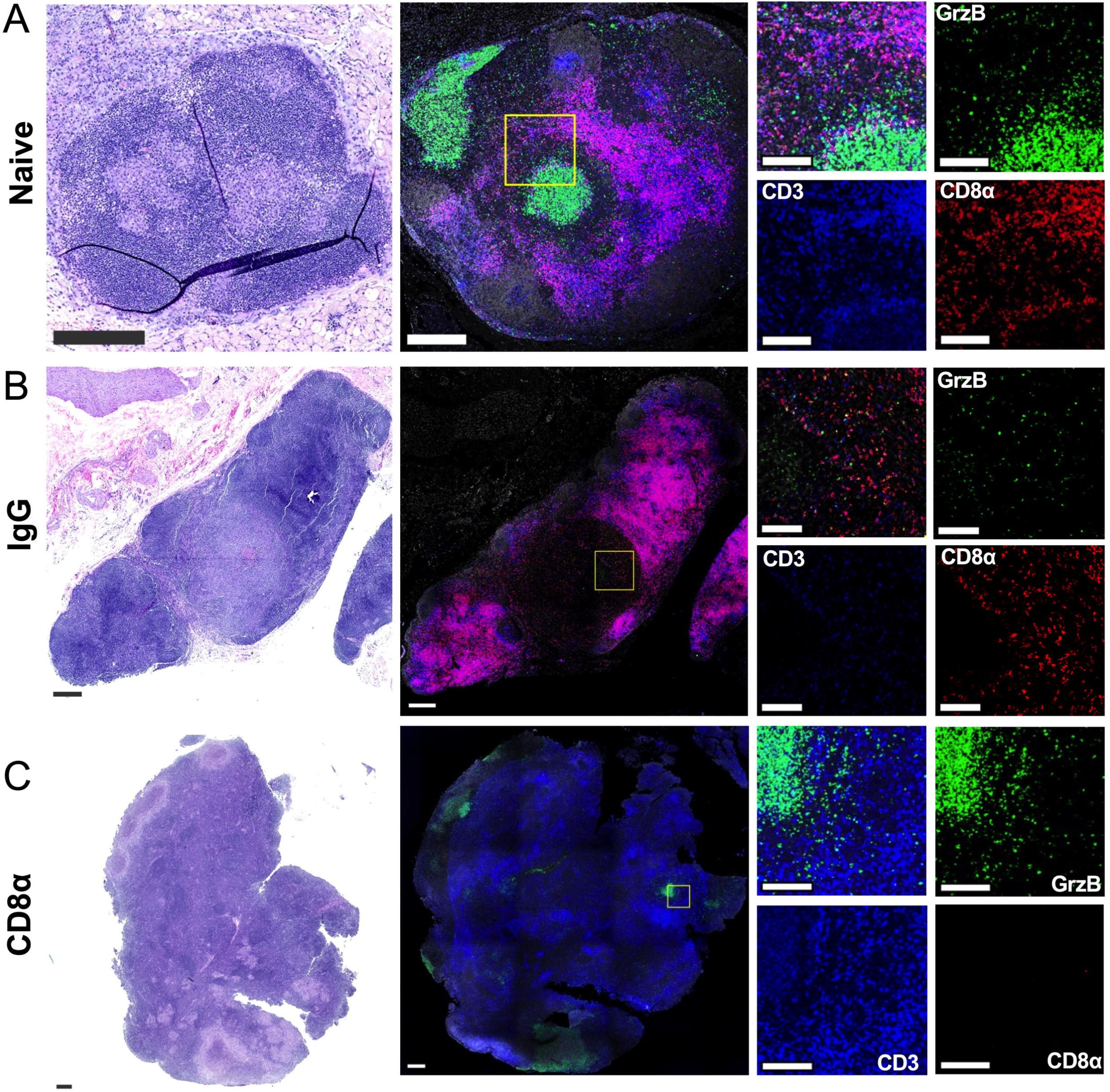
Localization and function of CD8α+ lymphocytes in LN granuloma. Select FFPE, involved thoracic LN from each cohort were stained with hematoxylin and eosin (H&E, *left* column) to characterize disease and tissue morphology followed by IFA staining (*middle*, *right* columns) with DAPI (*grey*), CD8α (*red*), Granzyme B (*green*) and CD3 (*blue*) to assess lymphocytes. (**A**) Left paratracheal LN from a naïve infected animal. (**B**) Left cranial hilar LN from an IgG-treated animal. (**C**) Left mainstem bronchus LN from a depleted animal. For all large images, scale bars represent 300μM. For all inset images, scale bars represent 100μM.

In some, but not all, CD8 depleted LNs, we observed small, organized clusters of CD8α+ cells (**Fig S6B**). Their morphology did not indicate a lymphocytic lineage, suggesting these cells may represent follicular dendritic cells (FDC). We stained serial sections from each cohort for CD20, CD21 (an FDC marker), and CD3; the CD8α+ clusters overlapped with CD21+CD20–CD3–staining. These regions were closely localized with a CD21+CD20+CD3–population, signifying follicular B cells, supporting that these CD8α+ cells are indeed FDC (**Fig. S6C**) and were not depleted with anti-CD8α Ab. Clusters of these CD8α+ cells can be identified by morphology and CD21 staining in both uninvolved and naive tissues, indicating that this is not an induced phenotype resulting from depletion or antibody infusions generally.

### Efficient CD8+ T cell cytotoxicity associates with contained lymph node infection

By flow cytometry, the relative abundance of myeloid populations is increased in involved (i.e. CFU+) LN tissues in both naïve and CD8α depleted reinfected animals (**Fig. 5A**). The lack of a noticeable change in IgG reinfection controls is likely a result of low bacterial burden even in involved LNs, resulting in contained lesions rather than effacement. The composition of lymphocyte subsets did not change across conditions, with the exception of expected losses of CD8+ cell types (**Fig. 5B**). Myeloid (p = 0.0665) and B cells (p = 0.1386) show a trend of a slight expansion by number per gram of tissue in involved LNs of naïve animals, which would be unsurprising given their role as professional antigen presenting cells and that these animals would be in the process of mounting an adaptive response to Mtb. (**Fig. 5C-M**). Functionally, conventional CD4+ and CD8+ T cells were not remarkably active. This is predictable because most cells in LNs are naïve, thus few activated cells reside there. Despite low frequencies in involved tissues, LNs in reinfected animals mount more efficient cytotoxic responses than those in naïve animals (**Fig. S7**). This efficient cytotoxicity may be a key functional aspect of CD8+ populations.

**Figure 5.**
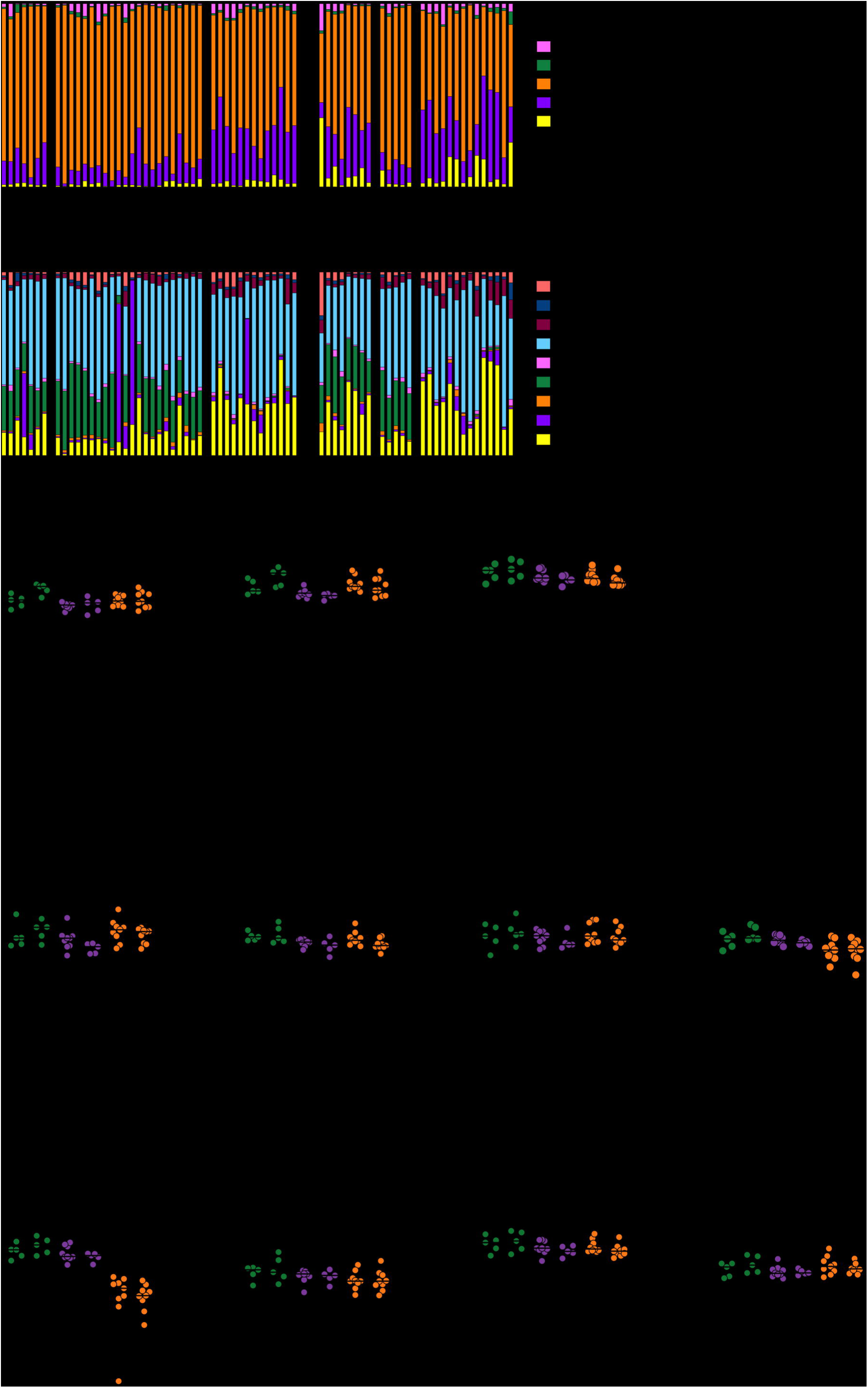
Composition of LN tissues. (**A-B**) Relative abundance of cell types in lymph nodes. Each bar represents an individual tissue. (**A**) Frequency of general immune populations as a percentage of live leukocytes. (**B**) Frequency of lymphocyte subsets as a percentage of all lymphocytes. For panels A-B, each bar represents a lymph node. (**C-L**) Number of (**C**) myeloid cells, (**D**) B cells, (**E**) total T cells, (**F**) γδ T cells, (**G**) NK cells, (**H**) ILCs, (**I**) CD8αα+ T cells, (**J**) CD8αβ T cells, (**K**) CD4+CD8+ T cells, (**L**) CD4+ T cells, (**M**) CD4-CD8-T cells per gram of tissue in uninvolved and involved LNs by depletion group. For panels C-M, p values shown are from paired t test comparing uninvolved and involved states within each treatment group. Each symbol represents an animal; line represents group median.

We could not establish a clear relationship between bacterial CFU in a LN and the number of barcodes detected (**Fig. S5D-F**). In fact, LNs with 1 barcode can reach similar peak bacterial burdens as those with 10 barcodes. Therefore, some level of protection against bacterial outgrowth conferred by concomitant immunity must occur upon establishment within LNs. The number of conventional CD8αβ T cells, the predominant CD8α+ cell type present in LNs, is negatively correlated with overall bacterial burden in LNs (**Fig. S5G**). Other metrics of LN involvement, such as overall unique barcodes, the number of CFU+ LN and LN bacterial burden do not seem to be related (**Fig. S5H-J**).

### Granuloma architecture is insufficient to contain Mtb in the absence of CD8+ lymphocytes

While less striking than the increase in Mtb burden within LNs, the partial loss of protection in the lung also suggests a role for CD8+ lymphocytes in containing infection prior to dissemination. Lung granulomas are typically organized into three key regions – (1) a core consisting by free bacteria and necrotic debris (i.e. caseum), (2) a surrounding myeloid region consisting of epithelioid and other macrophages, giant cells, dendritic cells and neutrophils, and (3) a lymphocyte cuff that encapsulates the lesion and isolates the site of infection from surrounding tissues. Recent spatial analysis of cellular organization in granulomas suggests that CD8+ T cells reside towards the outer edge of the lymphocyte cuff.^29^ The integrity of this organized structure contributes to the overall function and success of the lesion. We hypothesized that this peripheral distribution allows CD8+ populations to prevent bacterial outgrowth and escape. To query the impact of CD8α depletion on the spatial organization of lymphocytes within granulomas, we stained formalin-fixed paraffin embedded (FFPE) tissue sections for histologic analysis, by hematoxylin and eosin (H&E), and immunofluorescence (IF), specifically evaluating CD3, CD8α, GrzB, and CD4 localization.

Across treatment groups, typical lesion structure was generally conserved (**Fig. 6A-C**, **Fig. S8A-C**). The inner necrotic cores had characteristically low cellularity by DAPI (*white*) paired with GrzB staining (*green*) from activated cells that are constantly exposed to Mtb antigens. This core region was surrounded by presumed myeloid populations (CD3-), followed by a lymphocyte cuff with abundant CD3+ stained T cells (*blue*). Regardless of reinfection or depletion status, lymphocyte cuffs contained CD4+ T cells (**Fig. S8C**, *red*). In naïve primary lesions and IgG reinfection lesions, the cuff also contained the majority of CD8α+ cells (*red*) (**Fig. 6A,B**, **Fig. S8A**). However, this population was greatly reduced in the CD8α depleted group, supporting flow cytometric analysis (**Fig. 6C**, **Fig. S8B**).

**Figure 6.**
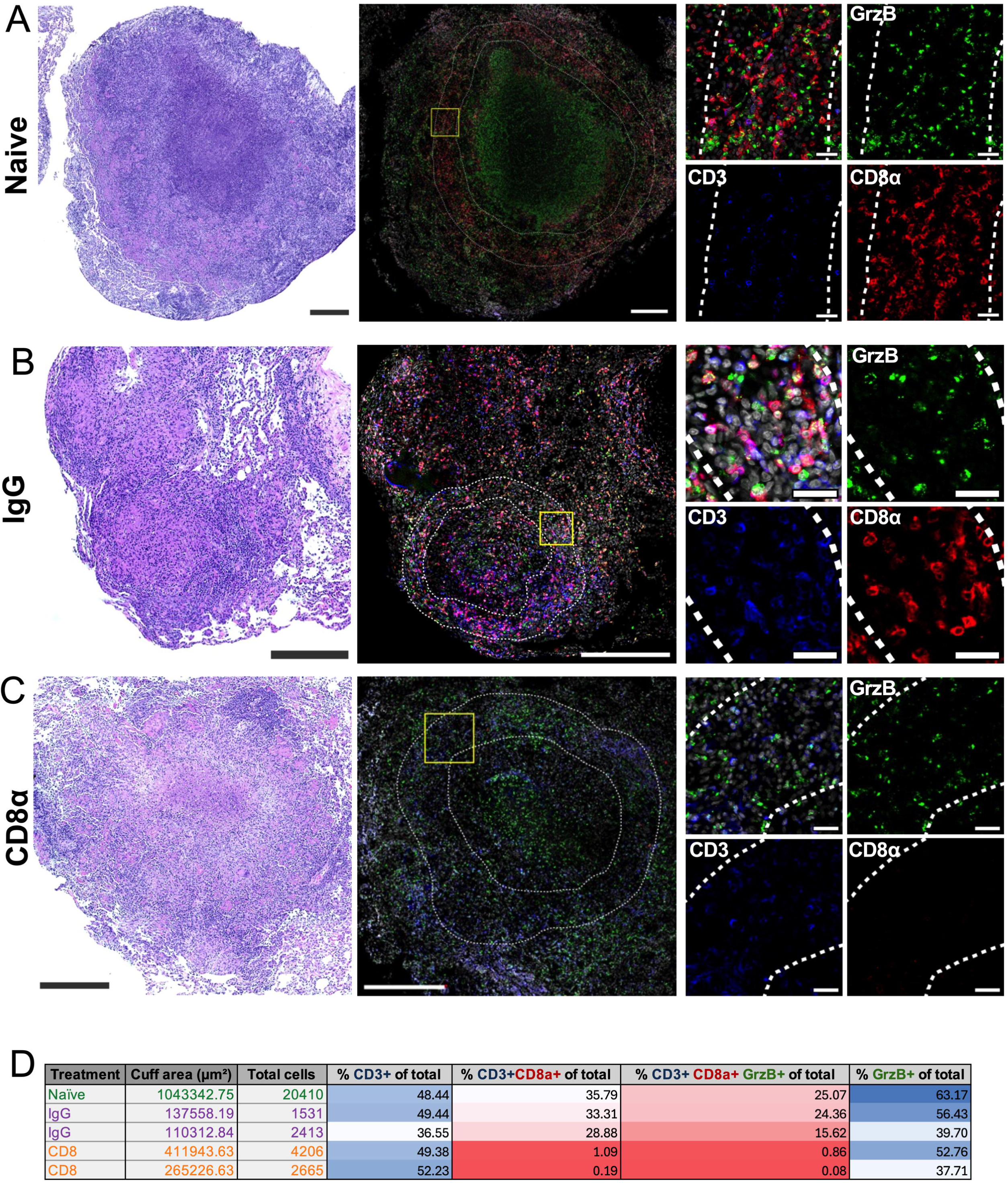
Localization and function of CD8α+ lymphocytes in lung granuloma. (**A-C**) FFPE lung granulomas from (**A**) a naïve animal, (**B**) an IgG animal, and (**C**) a depleted animal were stained with hematoxylin and eosin (H&E, *left* column) to characterize disease and tissue morphology followed by IFA staining (*middle*, *right* columns) with DAPI (*grey*), CD8α (*red*), Granzyme B (*green*) and CD3 (*blue*) to assess lymphocytes. For each image in panels A-C, scale bars for large images (*left*, *middle* columns) represents 300μM and inset image (*right* column) scale bars represent 30μM. The dashed regions signify the lymphocyte cuff. (**D**) Quantification of lymphocyte cuff area and cell count as well as percentages of CD3+ cells, CD8 α + T cells, Granzyme B+ CD8α+ T cells, and Granzyme B+ cells across each lung granuloma imaged (panels A-C, Fig. S7 panels A-B). For each column, blue represents higher values and red represents lower values where dark blue is 100% and red is 0%.

To quantify these characteristics, we further scrutinized the lymphocyte cuff by denoting a region of interest (ROI), informed by CD3+ staining. The total number of cells in this region varied across lesions but remained proportional to size (i.e. cross-sectional area of ROI). The frequency of CD3+ lymphocytes as a fraction of all cells detected was consistent across all tissues stained. This again highlights that successful protection conferred by concomitant immunity does not drastically alter the composition of cell types recruited to the site of infection, even if fewer cells are ultimately more effective at containing the bacteria. The level of CD3+CD8α+ T cells dropped in depleted tissue, where the percentage of CD8 T cells decreased from >30% to nearly 1% (**Fig. 6C-D, Fig. S8B**). Since the frequency of T cells detected within the ROI is consistent, this implies that there is still successful recruitment from surrounding tissues, rather than a loss of CD4+ cells within lesions.

Consequently, the relative expansion of the CD4 population in the absence of CD8+ T cells suggests compensation for missing roles in responding to increased disease burden. While a majority of CD8 T cells display GzmB+ signal, the level of GzmB+ in all lymphocyte cuff cells does not measurably drop. Similar results were seen following either CD8α or CD8β depletion in acute Mtb infection, where remaining CD4+ T cells compensated for some (e.g. granzymes), but not all (e.g. perforin), cytotoxic functions lost as a result of depletion.^8,18,29^

These results suggest that while general lesion organization and GzmB production remain following CD8α depletion, successful containment of Mtb relies on CD8+ cell types. Direct killing of extracellular pathogens by NK cells and infected cells by CD8+ T cells heavily relies on perforin and granulysin, which are generally lacking from CD4 T cells. Multiplexed ion beam imaging has demonstrated that the relative position of CD4 T cells is skewed towards the granuloma core relative to CD8 T cells.^29^ Thus, these results together suggest that close proximity interactions between CD4 T cells and the myeloid-dense core are needed to limit the bacterial burden, while CD8+ lymphocytes serve as peripheral sentries responsible for preventing escape (i.e. containment failure) and limiting overall bacterial growth.

### CD8+ and CD4+ cells control independent facets of bacterial outgrowth in granulomas

Unlike CD4 depletion, depletion of CD8α expressing cells did not lead to increased average CFU per granuloma (**Fig. 7A**). Thus, limiting bacterial growth (or increasing bacterial killing) within lesions is likely driven by CD4 T cell responses via interactions with myeloid populations, with CD8+ lymphocytes contributing a separate but equally critical role in infection outcome.

**Figure 7.**
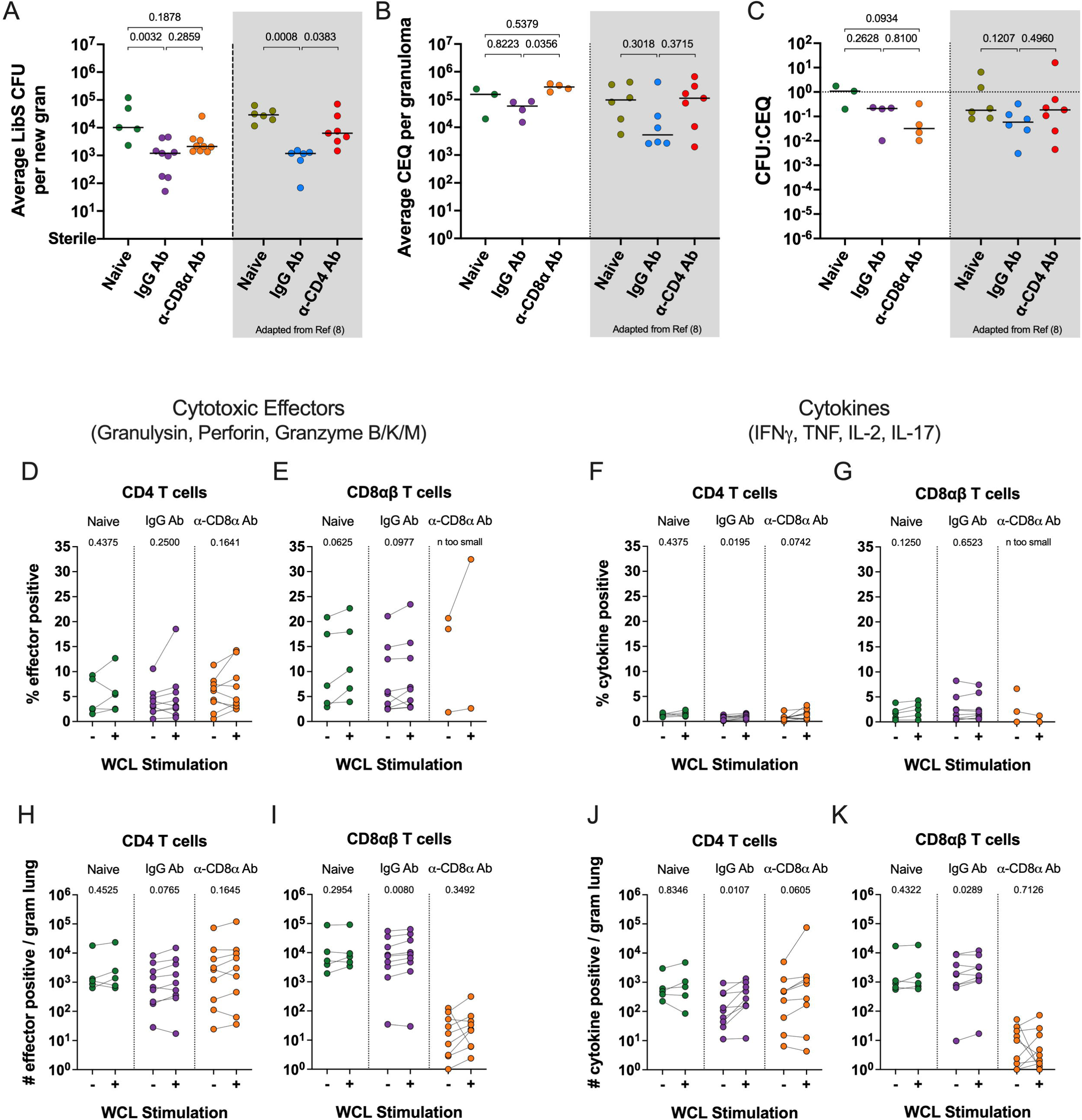
T cell function in lung tissue. (**A**) Mean bacterial burden per individual granuloma in each animal. (**B**) Mean number of chromosomal equivalents present in granulomas. (**C**) Killing capacity in granulomas, shown by the ratio between CFU and CEQ. For panels A-C, p values shown are for Welch’s ANOVA with Dunnett’s T3 multiple comparisons tests. In CD8α depletion study, all groups were compared against each other. In CD4 study, all groups were compared to the non-depleted IgG reinfection control. Each symbol represents an animal; line represents group median. (**D-G**) Percent of cells in lung tissue producing effector molecules with or without stimulation with Mtb whole cell lysate (WCL). P values shown (**H-K**) Number of cells per gram of lung tissue producing effector molecules with or without stimulation with WCL. Panels D,E,H,I show cytotoxic effector molecules; panels F,G,J,K show cytokines. Panels D,F,H,J show CD4 T cells; panels E,G,I,K show CD8αβ T cells. For panels D-K, stimulation conditions were compared within treatment groups. For panels D-G, p values shown are for Wilcoxon signed-rank non-parametric test. For panels H-K, p values shown are from paired t test. Each symbol represents an animal, lines connect animals across stimulation conditions.

While CFU serve as a single, and final, snapshot of granuloma containment, it does not fully convey the full dynamics of the host-pathogen interaction. For example, one granuloma with relatively static bacterial growth and negligible clearance by immune cells could have similar burden (by CFU) to another with constant bacterial growth combatted by efficient bacterial killing. Since bacterial genetic material remains within lesions for extended periods, we can extract and quantify chromosomal equivalents (CEQ) via PCR.^3,8,30^ In contrast to CFU, which enumerates the number of live bacteria present in the tissue at a specific moment, CEQ estimates the total number of bacteria to have ever been in the lesion. Here, all granulomas from a subset of animals were analyzed, covering the full range of CFU (**Fig. S9A**). Granulomas from CD8 depleted animals contained significantly higher CEQ than non-depleted reinfected controls, suggesting that bacterial killing still occurred within the lesion (**Fig. 7B**). The ratio of CFU to CEQ in each lesion indicates the level of bacterial killing that occurred, with no killing occurring when CFU and CEQ are equal (**Fig. 7C**). Evidence of substantial bacterial killing is seen consistently across animals and at the individual lesion level (**Fig. S9B,C**). When comparing the values within individual granulomas, granulomas in naïve animals exhibited poor ability to kill Mtb, indicated by little to no difference between CFU and CEQ levels, which is unsurprising only four weeks following initial exposure to mycobacterial pathogens with no established adaptive immune response (**Fig. S9B,C**). Both reinfected groups showed evidence of successful killing (ratio less than 1, **Fig. 7C, Fig. S9C**; positive slope from CFU to CEQ, **Fig. S9B**), although the magnitude of each metric was higher in depleted animals. Thus, these cell types are likely crucial mediators on preventing overwhelming outgrowth and, ultimately, lesion failure leading to dissemination.

### Cellular composition of LibS granulomas is altered by depletion

No significant differences were noted in granuloma size, number of cells, or inflammation as measured by PET CT between groups, which is a consistent observation across studies (**Fig. S9D-H**).^8^ We analyzed the composition of immune populations within lung granulomas by spectral flow cytometry (**Fig. S10**). The relative abundance of lymphocyte subsets was not noticeably changed by previous infection, further suggesting that enhanced protection is not mediated by altered recruitment of cell types (**Fig. S10A**). Following CD8α depletion, the frequency of CD8αα T cells, CD8αβ T cells, CD4+CD8α+ T cells, and NK cells decreased as expected. In contrast, reinfected animals had a significantly reduced number of B cells, and trends towards fewer conventional CD4 (p = 0.0544) and CD8αβ (p = 0.0889) T cells, in granulomas compared to naïve controls (**Fig. S10B-F**). It remains unclear whether this is a function of disease burden or a characteristic feature of protective lesions in concomitant immunity because there is little overlap between outcomes in the naïve and IgG groups. Similar consistency in lymphocyte composition is observed in uninvolved lung tissue (**Fig. S10H**). However, beyond the expected depletion of CD8α+ cell types, no significant changes were observed in the number of cells per gram of tissue (**Fig. S10I-M**). These data further support findings in other studies of concomitant immunity in TB that protection is not necessarily related to which cell types are present, but rather their functions that determine the effectiveness of protection.^31,32^

### Mycobacteria-specific responses persist in lungs of reinfected animals following drug treatment

Due to the small number of cells overall and each T cell subset specifically, especially in reinfection granulomas that are controlling Mtb, it is difficult to detect true functional responses using flow cytometry on granuloma homogenates. Further, Mtb-specific responses are difficult to quantify due to endogenous stimulation from bacteria, antigens and APCs within the lesion. However, in animals with prior Mtb infection, we reasoned that T resident memory cells in lungs were likely responsible for initial control of infection upon re-exposure. Thus, we analyzed T cells from enzymatically digested lung tissue, from which we could collect higher numbers of cells, for production of five cytotoxic effectors – granulysin, perforin, and granzymes (Grz) B/K/M – and four cytokines – IFN-γ, TNFα, IL-2, and IL-17 – in response to Mtb whole cell lysate (WCL; **Fig. 7D-K**). Since CD8αβ T cell depletion was very effective in lung tissue, the frequency data for this cell type in depleted animals (panels E and G) are not interpretable due to the low number of recorded events (see Methods).

The lack of antigen specific function in naïve animals is attributable to the early necropsy timepoint preempting an adaptive response mounted against mycobacteria, which can take longer than 6 weeks to mature. More T cells (CD4+ and CD8αβ) per gram of lung tissue in non-depleted reinfected animals produced cytotoxic effectors and cytokines in response to WCL stimulation (**Fig. 7H-K**). A similar trend (p = 0.0605) was seen in cytokine production, but not cytotoxic function, by CD4+ T cells in CD8α depleted animals (**Fig. 7H,J**). This indicates that following depletion of CD8+ cell types, critical immune functions are absent or diminished with minimal compensations by CD4 T cells. This effect, paired with high CEQ levels detected in depleted lesions, may be a reflection of ineffective “moonlighting” seen in CD4 T cells following depletion of CD8+ cell types (**Fig 7H**).^18^ In parallel, the infection worsens, further altering the immune state. This suggests that CD8+ cell types, at least in a natural model of immunity, are responsible for clearance/killing rather than controlling the pro/anti-inflammatory environment, which is a distinct and specialized role played by CD4s.

## DISCUSSION

Concomitant immunity leads to robust protection against subsequent Mtb exposure leading to active TB in humans and thus can provide clues to how best to design an effective vaccine. Our ability to model reinfection in a controllable NHP system allows us to gain valuable understanding into how humans in endemic regions respond to frequent exposures and elucidate immune mechanisms that can guide vaccine design. Previous work from our group has established that treatment of the initial infection partially abrogates protection against subsequent exposure and that CD4 T cells are necessary to maintain a protective response to subsequent exposures. The CD4 depletion reinfection study supported that CD4 T cells are critical in balancing the need for mounting a strong response (i.e. pro-inflammatory, driven by IFN-γ) and resolving the infection (i.e. anti-inflammatory, driven by CD4 modulation of immune responses leading to increased IL-10 and TGF-β). Further, the loss of CD4 T cells did not completely abrogate protection following reinfection but rather led to an intermediate state – more protected than immunologically naïve animals and less protected than animals with prior exposure to Mtb. These data suggest that other cell types are contributing to protection against TB. In the current study, we demonstrate that CD8α+ lymphocytes (innate-like and/or adaptive) are required for protection against reinfection, specifically in limiting dissemination to and growth within thoracic lymph nodes, a role distinct from CD4 T cells.

Here, we administered either a CD8α depleting antibody or an IgG isotype control antibody to macaques that had previously been Mtb-infected and treated with anti-TB drugs. We demonstrated successful depletion in the airways, lung and lymphoid tissues. Depletion of CD8α expressing cells resulted increased thoracic bacterial burden compared to IgG controls, demonstrating a clear role for CD8α+ cells in immunity against Mtb. Similarly, in a depletion study in IV-BCG vaccinated NHP, both CD4 T cells and CD8α+ lymphocytes were required for protection against TB.^17^ Thus, CD4 T cells alone cannot confer protective immunity against Mtb in these scenarios. Analysis of T cell functional profiles underlying these findings were consistent across flow cytometry, scRNAseq and immunofluorescence imaging, and aligned with fundamental findings in previous experiments. CD4+ and CD8α+, both innate and adaptive, populations act in parallel to provide protection against reinfection as well as protection provided by IV BCG, unable to fully compensate for the other.^8,17^ This has critical implications for vaccine strategies against TB, where simply inducing CD4 T cells is unlikely to be sufficient for full protection. This should guide strategies for designing, characterizing and evaluating future vaccine platforms to mount robust immune responses.

An emerging theme in the field is that Mtb infection can be contained at multiple stages and in distinct compartments. Here, we found that airway T cell populations are functionally altered by primary infection, with an increase in both CD4+ and CD8+ subsets producing cytokines and cytotoxic effector molecules, whether by expansion or recruitment. Transcriptomic analysis corroborated these findings, along with parallel shifts in the expression of associated transcription factor genes. Prior to secondary Mtb challenge, CD4 T cells in the airway of depleted animals adopted a partial, yet incomplete, cytotoxic profile in the absence of CD8+ subsets, mirroring ineffective moonlighting features observed in granulomas following CD8 depletion during primary Mtb infection.^18^ This reproducible phenomenon, found across mucosal surfaces and parenchymal lesions, suggests a mechanism by which T cell functional sensing and plasticity extends beyond helper phenotypes to missing cytotoxic responses. However, the limited breadth of these profiles, generally consisting of granzyme B without perforin or granulysin, appears to hinder the extent to which CD4 subsets can compensate for an insufficient contribution from the CD8+ compartment. Given the significant dissemination to lymph nodes observed in CD8α depleted animals, a competent cytotoxic response during establishment may be critical in preventing early escape from developing lesions.

Additionally, NK cells in the airway are associated with limiting initial successful establishment of Mtb. Therefore, NKs appear to be a consistent factor across successful models of protection, having been identified as an airway correlate of IV BCG mediated immunity, and in the blood of humans with LTBI.^24,33^ How to engage NK cells in a vaccine strategy should be considered, as well as the functions of these cells that contribute to immunity. The number of NK cells in the airway and IL-17 production by NK cells were noted in IV administration of BCG in macaques, suggesting a consistent immune mechanism involved in protection against Mtb spanning vaccination and natural immunity. These models of protection against Mtb infection have indicated that population level compositional shifts are not the sole driver of protective immunity; instead, temporal and functional dynamics are key to efficient responses. However, many analyses rely on a small number of functional markers, such as IFN-γ from CD4 T cells, for simplicity, which is unlikely to provide sufficient information about protective potential. Another consistent finding across our model of concomitant immunity and natural protection, generally, is an emergence of Granzyme K as a prevalent functional factor that is increased in protected groups. It remains unclear what specific role granzyme K plays, but this effector molecule and others warrant attention for evaluating host-directed therapy and vaccines.

A fundamental finding of this study was the significant and substantial increase in disseminated disease, specifically in thoracic lymph nodes, in CD8-depleted animals. Although it is not understood if dissemination occurs with free bacteria or those residing with phagocytes, such as dendritic cells or macrophages, LNs are a critical component in TB progression. The results of this study differ strikingly from the modest effects on extensive LN disease following CD4 depletion. Thus, we have demonstrated a unique role for CD8α+ lymphocytes in limiting dissemination of Mtb. During initial exposure and establishment of the infection, animals lacking CD8+ lymphocytes showed a slight increase in the number of barcodes established, which was also seen in CD4 depletion. In CD8-depleted animals, the number of dissemination events (unique barcodes reaching LN), the number of Mtb-infected LNs in each macaque, and bacterial burden in LNs increased, indicating that there may be multiple CD8 cell-related axes at play. LNs are a unique, immune privileged tissue with relatively little NK and innate immune activation. Further, there is no relationship between the number of barcodes and bacterial burden. Together, this implicates adaptive cell types in responses to control outgrowth once bacteria reach the LN. The lack of bacterial control in the LN indicates that CD8+ populations have a role in limiting replication or even killing Mtb in lymphoid tissues. In a previous study by our group, CD8α and CD8β expressing cells were depleted via antibody infusion prior to challenge with Mtb in naïve macaques. Both depletion groups showed significant increases in thoracic LN CFU.^18^ Histologically, the loss of CD8α+ lymphocytes in the LN results in pathology reflective of naïve LN, where infection disrupts CD3+ lymphocyte organization and results in multifocal lesions. Given that the majority of CD8α+ cells in LNs are CD8αβ T cells, it is highly plausible that functional CD8αβ T cells producing GzmB and other cytotoxic effector molecules are a critical component of LN protection while a broader population of CD8α+ lymphocytes, including innate-like cells, contribute to limiting bacterial shuttling from the lung. While LN dissemination increased in the context of reinfection in the absence of CD4 T cells, effects were most prevalent within the lung and bacterial burden remained lower in distal sites of infection.

Recent work by McCaffrey et al demonstrated spatial skewing of CD4 T cells towards the myeloid core and CD8 T cell populations towards the periphery.^29^ Our evaluation of CFU and CEQ implies that CD8+ lymphocytes may be involved in preventing a permissive growth environment (i.e. high CEQ while CFU remains similar to IgG in depleted animals). CD4 T cells are also important in limiting growth in granulomas but are primarily critical in the context of killing Mtb (high CEQ with high CFU in depleted animals compared to IgG). A loss of containment within the lung and dissemination to the LNs supports that the CD8+ cells residing within the lymphocyte cuff may act as “sentries”, maintaining lesion integrity and preventing or limiting escape of Mtb from the granuloma. This also supports that CD8+ lymphocytes strongly contribute to the compartmentalization of TB disease, separately affecting initial establishment, granuloma bacterial growth, dissemination to, and control within the thoracic LNs.

CD8α depletion can also affect a subset of dendritic cells, most prominently conventional type 1 dendritic cells (cDC1). This population is a key contributor in shuttling Mtb from the lungs to LN, initiating antigen specific memory responses, which is a normal phenomenon in Mtb infection. However, in our model, an adaptive response would have formed prior to depletion, so memory populations have been established. Single cell RNA sequencing analysis in a previous study by our group showed no change in cDC1 or cDC2 populations following CD8α depletion, suggesting this is a durable population unaffected by our experimental interventions.^18^

The conclusions of this study are primarily limited by an inability to determine the specific CD8+ lymphocyte subsets that are playing a role in protection. With CD8α depletion, a broad spectrum of immune cell types is affected across the innate-adaptive spectrum, including conventional CD8 T cells (CD8αβ T cells), unconventional or innate-like T cells (e.g. MAITs, NKTs, and some γδ T cells), CD4+CD8+ T cells, and NK cells. In future studies, an anti-CD8β antibody could be used to more specifically deplete adaptive CD8 T cells to determine the role of this adaptive immune subset, as we have published in our IV-BCG vaccine model. At this time, reagents do not exist that can reliably and specifically target the innate-like CD8+ lymphocytes for depletion; this remains a goal for the future. We used imaging, detailed necropsy protocols and barcoded strains of Mtb to quantify LN involvement. However, defining and untangling protective mechanisms specific to lung or LN compartments may require ex vivo, explant and organoid models of infection. Lastly, when we implement antibody mediated depletion in our model of reinfection in macaques, it necessitates antibiotic treatment of the primary infection to prevent rapid progression even prior to a second challenge. Elimination of the primary infection likely limits the amount of persistent antigen remaining in the lung which is needed to maintain T resident memory cells. However, the consistent treatment of both reinfection groups (i.e. depletion and immunocompetent), along with similar outcomes across multiple historical studies, provides a reliable baseline from which to form conclusions.

These results also highlight that the mechanisms of protection or bacterial control are likely different in unique models of protection against TB. The data presented here demonstrate that, in the context of concomitant immunity, some CD8+ cell types, both innate and adaptive, are critical during the earliest interactions with the pathogen, as well as at later timepoints as disease progresses. Our previous data support that CD4 T cells have multiple roles in protection against reinfection, but one of those roles seems to be an immunoregulatory role affecting CD8 T cells.^8^ CD8+ lymphocytes are also critical for limiting acute infection, as well.^18^ Both CD4 T cells and innate-like CD8+ lymphocytes are necessary for the robust protection induced by intravenous BCG in macaques, and a previous study indicated a role for CD8+ lymphocytes in BCG ID induced protection.^17,34^ Thus, successful vaccine strategies likely depend on a wide range of immune lymphocytes and functional responses, as well as positioning those cells in the tissues to immediately limit Mtb replication.

## ACKNOWLEDGEMENTS

We would like to acknowledge Joshua T Mattila (University of Pittsburgh School of Public Health) for providing technical expertise and imaging facilities and Jacob Klein (University of Pittsburgh School of Medicine) and Charlie Demurjian (MIT BioMicro Center) for their efforts towards making data publicly accessible. We are extremely grateful to the research technicians in the Flynn lab for their outstanding support of this work, and to the entire TB Research Group at University of Pittsburgh for advice and insights over the course of this study. This work was supported by the National Institutes of Health (NIH) National Institute of Allergy and Infectious Diseases (NIAID) (75N93019C00071 [IMPAcTB / HI-IMPACT], 5T32AI089443, and 5T32AI060525). Operations of The University of Pittsburgh Regional Biocontainment Laboratory (RBL) within the Center for Vaccine Research (CVR) are supported by the NIH NIAID (UC7AI180311).

## AUTHOR CONTRIBUTIONS

Conceptualization: SMF, JLF; Data curation: AWS, MCC, RJK, CLA, PM; Formal analysis: AWS, MCC, GB, MRC, LEH, PM; Funding acquisition: SMF, JLF; Investigation: AWS, GB, MAR, LEH, FH, MRC, AC, AAS, RJK, CLA, EK, HC, MM, HJB, PM, PLL; Methodology: AWS, MCC, GB, FS, PM, JLF; Project administration: MAR, CAS; Resources: AKS, SMF, JLF; Supervision: MAR, PM, CAS, SMF, JLF; Visualization: AWS, MCC, GB, LEH, MM, PM; Writing – original draft: AWS, GB, JLF; Writing – review & editing: AWS, MCC, FS, AKS, SMF, JLF.

## DATA AND MATERIALS AVAILABILITY

All amplicon sequencing data in this work is publicly available on SRA under the accession number: PRJNA1513074. All primary data, including PET CT scans, are available publicly at: https://fairdomhub.org/studies/1515. Additional information is available from Dr. JoAnne Flynn upon request.

## METHODS AND MATERIALS

### Animals and Husbandry

Twenty-three (23) Chinese cynomolgus macaques (*Macaca fascicularis*; CCM) were obtained from Valley Biosystems. Due to limited availability of female macaques, this study consisted of 18 males and 5 females, aged between 4 and 12 years old (mean 8.4). All work involving CCMs was conducted at the University of Pittsburgh School of Medicine and procedures complied with Institutional Animal Care and Use Committee ethical regulations. All animals were housed and cared for in accordance with local, state, federal, and institute policies in facilities accredited by the American Association for Accreditation of Laboratory Animal Care, under standards established in the Animal Welfare Act and the Guide for the Care and Use of Laboratory Animals as mandated by the U.S. Public Health Service Policy. Upon arrival at the institution, CCMs were quarantined and evaluated by interferon-gamma release assay (i.e. IGRA) to determine evidence of pre-existing mycobacterial infection. Signs of general wellness such as behavioral changes, body condition, temperature, and blood panels were monitored throughout the study. Following Mtb challenge, veterinary staff monitored animals for clinical and bacteriologic signs of disease, including changes to appetite, weight, activity levels, coughing, erythrocyte sedimentation rate (ESR) and positive gastric aspirate cultures. Paired assessment of PET CT imaging with these metrics determined whether an animal met humane endpoint criteria prior to the scheduled endpoint of the study.

### NHP Study Design

Eighteen (18) CCMs were randomized into two groups (**Table S1**). Two groups (n = 9 each) were infected with Mtb (barcoded Erdman strain, Library P; LibP) for nine weeks before initiation of antibiotic therapy with a standard regimen (isoniazid (H), rifampin (R), pyrazinamide (Z), ethambutol (E); HRZE). Two animals had rapidly progressing disease prior to the scheduled initiation of HRZE and were subjected to additional monitoring to ensure proper care, with one starting antibiotics ∼ 3 weeks early. These two animals were separated experimentally, with one being put in each arm (IgG and anti-CD8α). After 18 weeks of treatment, CCMs were rested for approximately five weeks prior to a secondary, 4-week Mtb infection (barcoded Erdman strain, Library S; LibS). Two weeks prior to the second challenge, these CCMs began receiving intravenous antibody infusions, which continued every two weeks until necropsy. A third randomized group (n = 5) was infected with LibS in parallel with the other groups as a naïve (i.e. no known previous mycobacterial exposure) control. This study was completed across two cohorts (n = 12, 11; **Table S1**).

### Bronchoalveolar Lavage

Bronchoalveolar lavage (BAL) were performed, as previously described, at four time points throughout the study – prior to LibP infection, following drug treatment of the LibP infection, and following depletion antibody infusion.^35^ Throughout the procedure, the breathing pattern and O_2_ saturation levels of animals were monitored by veterinary technicians. Animals were fasted overnight prior to the procedure, sedated with ketamine or telazol, pre-treated with 100% oxygen and positioned in dorsal recumbency. Lidocaine was also used to aid bronchoscope placement reduce sneeze/cough responses. A flexible pediatric bronchoscope was advanced past the mainstem bronchus, wedged into a distal subsegmental bronchus, and four 10 mL aliquots of warmed sterile saline were sequentially instilled and aspirated via the suction channel. Following the procedure, animals received 100% O_2_ via face mask for 5 min before being returned to their home cage and were monitored closely during recovery. Recovered samples were centrifuged to isolate cells and supernatant. Cells were resuspended in warm R10 media for immunologic assays. Remaining cells and fluid were cryopreserved for future analysis.

### Single cell RNA sequencing and genome alignment

Single-cell transcriptomes of BAL cells were generated with the Seq-Well S^3^ platform, as previously described.^36,37^ Briefly, cells were loaded onto Seq-Well arrays preloaded with uniquely barcoded mRNA capture beads and sealed with a semi-permeable polycarbonate membrane, followed by cell lysis, transcript hybridization, and bead recovery. Captured mRNA was reverse transcribed (Maxima H Minus Reverse Transcriptase, Thermo Fisher Scientific), excess primers were removed by exonuclease I digestion (New England Biolabs), and second-strand synthesis and whole transcriptome amplification (KAPA HiFi HotStart ReadyMix, Kapa Biosystems) were performed. Dual-indexed 3′ digital gene expression libraries were prepared with the Nextera XT kit (Illumina) and sequenced on an Illumina NovaSeq 6000 (S4 flow cells) at the Broad Institute, using a custom Read 1 primer and a paired-end read structure of Read 1, 21 bases (12-base cell barcode and 8-base unique molecular identifier, UMI); Index 1 and Index 2, 8 bases each; and Read 2, 50 bases (cDNA).

Reads were processed with the Drop-seq tools pipeline,^38^ as implemented in the Cumulus *dropseq_workflow* on the Terra platform (app.terra.bio).^39^ Each cDNA read (Read 2) was tagged with its cell barcode and UMI from Read 1, trimmed of template-switching oligonucleotide and poly(A) sequence, and corrected for bead synthesis and substitution errors. Reads were then aligned with STAR (v2.7.0c)^40^ to the *Macaca fascicularis* 5.0 genome with its NCBI RefSeq annotation (GCF_000364345.1).

### Transcriptional data processing

For individual samples, ambient RNA contaminating cell transcriptomes was estimated via decontX. Resulting gene expression matrices were filtered on the number of features (200 < x < 7500) and counts (300 < x < 50000). Further filtering removed cells with mitochondrial, ribosomal and beta-globin gene expression frequency above thresholds of 0.1 and likely doublets were identified with DoubletFinder. Initial clustering indicated batch effects from sample sequencing run, or study cohort that were addressed via Harmony integration. Coarse level clustering and cell type annotation was then performed to identify general populations, from which T- and NK-like cells could be isolated. The top 23 principal components (PCs) were used for dimensionality reduction with resolution set to 1.0, leading to the identification of 26 distinct clusters. Iterative subclustering on T/NK cells removed contaminants, primarily myeloid in nature, not identified during preliminary quality control processing, corroborated by parallel manual and automated referencing via ProjecTILs.^41^ High-quality T cells were then annotated by CD4-, CD8-, and NK-like transcriptional profiles. No statistical analyses were performed based on the number of cells and samples analyzed.

### Mtb Challenge

CCM were intrabrochially instilled with Mtb (Erdman strain), as previously described.^42,43^ Genetically barcoded libraries (LibP and LibS) were used to afford differentiation of lesion age and bacterial burden across multiple infections.^6^ Infection procedures were carried out by trained veterinary technician staff members with a target inoculation dose of 15 CFU. Dose was ultimately determined by plating, with a range of 17-32 CFU for LibP infections and 9-28 CFU for LibS CFU, which has resulted in successful establishment of Mtb infection in NHP previously (**Table S1**). PET CT scans 4 weeks post infection and barcoding analysis at necropsy corroborate these values.

### PET CT Scanning

PET CT imaging was performed using radiolabeled 2-deoxy-2-[^18^F]fluoro-D-glucose (FDG) in order to identify TB-associated lung lesions. Scans were obtained using a MultiScan LFER-150 PET CT scanner (Mediso Medical Imaging Systems) as previously described.^7,44,45^ Analysis of imaging focused on identification and enumeration of emergent lesions immediately prior to necropsy. Additional radiologic metrics include quantification and three-dimensional visualization of total pulmonary PET uptake exclusive of thoracic lymph nodes (Total FDG Lung activity), maximum PET uptake for each affected thoracic lymph node, as well as lesion size and PET uptake for each identified lung lesion. LNs do not readily appear on CT scans until they are enlarged due to infection, so only animals with observable enlarged and FDG-avid LNs could be analyzed by PET CT for LN related metrics.

### Necropsy and Tissue Processing

Approximately four weeks after LibS challenge, animals were terminally bled and euthanized by sodium pentobarbital injection, as previously described.^23^ The most recent PET CT scan was used to map known lesions and pathology, and gross pathology observed during prosection was quantified using a published scoring system.^23,43^ Regions of uninvolved tissue from each lung lobe, peripheral (e.g. axillary, inguinal) and thoracic (e.g. carinal, cranial hilar) lymph nodes, scan matched TB pathology (e.g. granulomas, consolidations, pneumonia) and any potential pathology identified upon gross examination of tissue were excised and processed for various immunologic and microbiologic analyses. Lung lobes were enzymatically dissociated (Gentle MACS; Miltenyi) and all other samples were homogenized by mechanical disruption in PBS. All single cell suspensions were filtered with a 70-um cell strainer. Portions of all tissues, excluding granulomas < 2 mm in diameter, were also formalin fixed and paraffin embedded for histological analysis.

### CFU Determination

Tissue homogenates were plated on 7H11 with or without PANTA antibiotic supplantation for quantification of bacterial burden. Colony forming units (CFU) were counted after incubation at 37°C in 5% CO_2_ for 21 days. Lesions and other TB related pathology detected by PET CT after LibS challenge were attributed to LibS if LibP barcodes were not detected during sequencing. Disease not discernable on scans were only attributed to LibS if LibS was detected and LibP was not.

### CEQ Quantification

At the time of necropsy, an aliquot of every tissue homogenate was frozen at −80°C for DNA extraction and CEQ analysis. Genomic DNA was extracted with a phenol-chloroform method previously described.^30,46^ In brief, samples were resuspended in Tris-EDTA buffer, pH 8.0 with 70°C UltraPure phenol:chloroform:isoamyl alcohol (25:24:1) and 0.1 mm zirconia-silica beads. Volumes were recorded throughout the process to normalize for loss. After a 10 min incubation, samples were vortexed twice for 5 minutes, with a 30 sec break, to disrupt the bacterial walls. Genomic DNA was removed in the aqueous layer after centrifugation. The phenol:chloroform:isoamyl alcohol extraction was repeated followed by a 30 min incubation at room temperature. DNA was precipitated with isopropanol and 3M sodium acetate and washed with 70% ethanol. Pellets were resuspended in nuclease free water and DNA concentration was determined using a spectrophotometer.

Quantification of chromosomal equivalents (CEQ) of *M. tuberculosis* was determined using real-time PCR of a single copy of the gene, sigF, as previously described.^30^ Primer and probes were purchased as a pre-mix PrimeTime qPCR assay (Integrated DNA Technologies) and the sequences are as follows, 5’3’: probe–FAM-TCG GAC TTC GTC TCC TTC-Iowa Black, sigF Fwd–GCG GGT CGG GCT GGT CAA C, and sigF Rvs–CCT CGC CCA TGA TGG TAG GAA C. Real time PCR was performed in duplicate for every sample. CEQ was quantified in relation to a standard curve of serially diluted known M.tb genomic DNA extracted from our stock Erdman M.tb. The ratio of CEQ and the live CFU from plating reflects the bacterial killing or growth in each sample.^30^

### Barcode Sequencing

To track the establishment and dissemination of Mtb, barcoded Mtb Erdman were plated from tissue homogenates and bacterial genomic DNA extracted, as previously described.^6^ Then, two sequencing strategies were used for barcode determination. First, all genomic DNA samples were subjected to transposon-mediated fragmentation (Nextera XT, Illumina) and whole genome sequencing on a NovaSeq S4 platform (150 bp, paired end sequencing). Barcodes were enumerated from the fastq files, as previously described in work from our group, which briefly involved motif finding to detect constant regions around the barcode sequences and quantitate unique barcodes associated with both LibP and LibS q-tags.^5^ All singleton barcodes were removed as sequencing artifacts. These scripts can be found in the following Github link: https://github.com/Fortune-Lab/Mtb-WGS-barcoding.

Second, samples that failed whole genome sequencing or yielded <10 barcode counts after quantitation, were then subjected to amplicon-based barcode amplification and sequencing to increase our sensitivity of detection. Fastq QC and barcode detection was performed as previously described, which includes motif detection and quantitative detection of true barcodes through the BARTI pipeline.^18^ These amplicon-based detection scripts can be found at the following Github link: https://github.com/sarahfortunelab/barcodetracking. All barcodes from both rounds of sequencing were combined for statistical analysis (Supplementary Table S2).

### Spectral Flow Cytometry

Single cell suspensions from tissue homogenates of excised lung lobe, lymph nodes and granulomas were stimulated and stained with fluorophore-labeled antibodies for phenotypic and functional characterization, as described previously.^47^ Cells were incubated at 37°C in 5% CO_2_ for 2 hours in R10 media. Samples were stimulated with a pool of Mtb H37Rv strain whole cell lysate (WCL) peptides (BEI Resources) at 1 μg/mL. Media alone and P+I served as negative and positive control conditions, respectively. GolgiPlug (BD Biosciences) was added to a final concentration of 10 μg/mL for 4 additional hours. Samples were stained with a fixable viability dye and antibodies targeting characteristic surface markers before fixation and permeabilization, allowing for subsequent intracellular cytokine staining following standard protocols. Panels are detailed in **Table S4**. Data collection was performed using a five laser Aurora on SpectroFlo software (16UV-16V-14B-10YG-8R; Cytek) at the University of Pittsburgh School of Medicine Department of Immunology’s Unified Flow Core. Analysis was performed using FlowJo software (v10.10.0; **Fig. S11**). Frequency analyses excluded cell types and associated functions with fewer than 100 total lymphocyte events or 50 events from the parent cell type to avoid skewing.^17^

### Histopathological and Immunofluorescence Analysis

Tissues were collected at necropsy, including randomly selected uninvolved lung and both thoracic and peripheral lymph nodes. Granulomas were prioritized for homogenization, but lesions large enough to bisect by scalpel were split between standard processing and fixation. Samples designated for histologic analysis were placed in 10% buffered formalin fixative, embedded in paraffin and sectioned into 5-10 µm thick slices by microtome. This sample preparation was performed by University of Pittsburgh histology core facilities.

For histopathological analysis, slides were stained with hematoxylin and eosin (H&E) using standard protocols and microscopically evaluated by an experienced pathologist.

For immunofluorescence, slides were deparaffinized in xylene and 95% ethanol and placed in a pH 9.0 buffer containing 20mM Tris/820µM EDTA/0.0001% Tween-20 to achieve antigen retrieval with a Retriever (Electron Microscopy Sciences). The sections were incubated for 30 min at room temperature with 1% BSA in PBS followed by overnight staining at 4°C with primary antibodies diluted in blocking reagent. Slides were washed 3 times in 1X PBS and incubated for 1 hour at RT with secondary species-specific antibodies diluted in blocking buffer. Slides were washed 3 times in 1X PBS and incubated with TrueVIEW Autofluorescence Quenching Kit (Vector) for 5 min at RT. Slides were washed an additional 3 times in 1X PBS and mounted with coverslips using ProLong Gold mounting media containing DAPI (Thermo Fisher Scientific). Slides cured for 24 hours at RT and were stored at 20°C until imaging on a Nikon Ti2 Eclipse confocal microscope (Nikon, Melville, NY). Staining panels are detailed in **Table S4.**

### Statistical Methods

All statistical tests were performed in GraphPad Prism (for macOS, Version 10.6.1). Shapiro-Wilk normality test was used to assess normality. When comparing means (or medians) of two independent groups, unpaired t test (with Welch’s correction for groups with different variances) was used for normally distributed data. The Mann-Whitney rank sum test was used as a nonparametric alternative. For comparing more than two groups, Welch’s ANOVA test with Dunnett’s T3 multiple comparison tests were used to compare pairwise groups with normally distributed data. Otherwise, Kruskal-Wallis test with Dunn’s multiple comparisons were used. For normally distributed measurements that were repeated, pairwise mean differences were assessed using paired t test for two groups and repeated measures one-way ANOVA with Dunnett’s multiple comparisons tests for more than two groups. Wilcoxon matched-pairs signed rank test was used to compare two group medians for data which did not follow normal distribution. Spearman’s rho (ρ) was used to assess the correlation between two measurements.

## Supplementary Materials

Figs. S1 to S11

Tables S1 to S4

## References

1. Joslyn, L.R., Flynn, J.L., Kirschner, D.E., and Linderman, J.J. (2022). Concomitant immunity to M. tuberculosis infection. Sci Rep 12, 20731. 10.1038/s41598-022-24516-8.

2. Andrews, J.R., Noubary, F., Walensky, R.P., Cerda, R., Losina, E., and Horsburgh, C.R. (2012). Risk of Progression to Active Tuberculosis Following Reinfection With Mycobacterium tuberculosis. Clinical Infectious Diseases 54, 784–791. 10.1093/cid/cir951.

3. Cadena, A.M., Hopkins, F.F., Maiello, P., Carey, A.F., Wong, E.A., Martin, C.J., Gideon, H.P., DiFazio, R.M., Andersen, P., Lin, P.L., et al. (2018). Concurrent infection with Mycobacterium tuberculosis confers robust protection against secondary infection in macaques. PLoS Pathog 14, e1007305. 10.1371/journal.ppat.1007305.

4. Ganchua, S.K., Maiello, P., Chao, M., Hopkins, F., Mugahid, D., Lin, P.L., Fortune, S.M., and Flynn, J.L. (2024). Antibiotic treatment modestly reduces protection against *Mycobacterium tuberculosis* reinfection in macaques. Infection and Immunity 92. 10.1128/iai.00535-23.

5. Simonson, A.W., Chao, M.C., Hood, L.E., Donlan, R.A., Hopkins, F., Chase, M.R., Vickers, A.J., Callendrello, A., Klein, E., Borish, H., et al. (2025). Protection against reinfection with Mycobacterium tuberculosis extends across heterologous Mtb lineages. Mucosal Immunol. 10.1016/j.mucimm.2025.10.010.

6. Martin, C.J., Cadena, A.M., Leung, V.W., Lin, P.L., Maiello, P., Hicks, N., Chase, M.R., Flynn, J.L., and Fortune, S.M. (2017). Digitally Barcoding Mycobacterium tuberculosis Reveals In Vivo Infection Dynamics in the Macaque Model of Tuberculosis. mBio 8. 10.1128/mBio.00312-17.

7. White, A.G., Maiello, P., Coleman, M.T., Tomko, J., Frye, L.J., Scanga, C.A., Lin, P.L., and Flynn, J.L. (2017). Analysis of 18FDG PET/CT Imaging as a Tool for Studying *Mycobacterium tuberculosis* Infection and Treatment in Non-human Primates. JoVE Immunologu and Infection, 56375. 10.3791/56375.

8. Bromley, J.D., Ganchua, S.K.C., Nyquist, S.K., Maiello, P., Chao, M., Borish, H.J., Rodgers, M., Tomko, J., Kracinovsky, K., Mugahid, D., et al. (2024). CD4+ T cells re-wire granuloma cellularity and regulatory networks to promote immunomodulation following Mtb reinfection. Immunity 57, 2380–2398.e2386. 10.1016/j.immuni.2024.08.002.

9. Bromley, J.D., Simonson, A.W., Shalek, A.K., and Flynn, J.L. (2025). Host-intrinsic and host-extrinsic factors modulate immunity to Mtb infection, reinfection, and noncanonical vaccination routes. Cell Rep Med 6, 102286. 10.1016/j.xcrm.2025.102286.

10. Sia, J.K., and Rengarajan, J. (2019). Immunology of Mycobacterium tuberculosis Infections. Microbiol Spectr 7. 10.1128/microbiolspec.GPP3-0022-2018.

11. Rijnink, W.F., Ottenhoff, T.H.M., and Joosten, S.A. (2021). B-Cells and Antibodies as Contributors to Effector Immune Responses in Tuberculosis. Front Immunol 12, 640168. 10.3389/fimmu.2021.640168.

12. Sankar, P., and Mishra, B.B. (2023). Early innate cell interactions with Mycobacterium tuberculosis in protection and pathology of tuberculosis. Front Immunol 14, 1260859. 10.3389/fimmu.2023.1260859.

13. Flynn, J.L., and Chan, J. (2022). Immune cell interactions in tuberculosis. Cell 185, 4682–4702. 10.1016/j.cell.2022.10.025.

14. Behar, S.M. (2013). Antigen-specific CD8(+) T cells and protective immunity to tuberculosis. Adv Exp Med Biol 783, 141–163. 10.1007/978-1-4614-6111-1_8.

15. Li, J., Simonson, A.W., and Flynn, J.L. (2025). CD8+ lymphocytes in immunity against tuberculosis. J Immunol. 10.1093/jimmun/vkaf268.

16. Bjornson-Hooper, Z.B., Fragiadakis, G.K., Spitzer, M.H., Chen, H., Madhireddy, D., Hu, K., Lundsten, K., McIlwain, D.R., and Nolan, G.P. (2022). A Comprehensive Atlas of Immunological Differences Between Humans, Mice, and Non-Human Primates. Front Immunol 13, 867015. 10.3389/fimmu.2022.867015.

17. Simonson, A.W., Zeppa, J.J., Bucsan, A.N., Chao, M.C., Pokkali, S., Hopkins, F., Chase, M.R., Vickers, A.J., Sutton, M.S., Winchell, C.G., et al. (2025). Intravenous BCG-mediated protection against tuberculosis requires CD4+ T cells and CD8α+ lymphocytes. J Exp Med 222. 10.1084/jem.20241571.

18. Winchell, C.G., Nyquist, S.K., Chao, M.C., Maiello, P., Myers, A.J., Hopkins, F., Chase, M., Gideon, H.P., Patel, K.V., Bromley, J.D., et al. (2023). CD8+ lymphocytes are critical for early control of tuberculosis in macaques. Journal of Experimental Medicine 220, e20230707. 10.1084/jem.20230707.

19. Selwyn, P.A., Hartel, D., Lewis, V.A., Schoenbaum, E.E., Vermund, S.H., Klein, R.S., Walker, A.T., and Friedland, G.H. (1989). A prospective study of the risk of tuberculosis among intravenous drug users with human immunodeficiency virus infection. N Engl J Med 320, 545–550. 10.1056/NEJM198903023200901.

20. Grant, N.L., Kelly, K., Maiello, P., Abbott, H., O’Connor, S., Lin, P.L., Scanga, C.A., and Flynn, J.L. (2023). Mycobacterium tuberculosis-Specific CD4 T Cells Expressing Transcription Factors T-Bet or RORγT Associate with Bacterial Control in Granulomas. mBio 14, e0047723. 10.1128/mbio.00477-23.

21. Nelson, C.E., Kauffman, K.D., Osum, K.C., Sakai, S., Buchanan, J., Chanchu, J.M., Cohen, M., Laux, J., Dougi, I., Barrows, K.M., et al. (2025). Mycobacterium tuberculosis peptide-specific T cells in pulmonary granulomas display broad effector functions. iScience 28, 114034. 10.1016/j.isci.2025.114034.

22. Peters, J.M., Irvine, E.B., Makatsa, M.S., Rosenberg, J.M., Wadsworth, M.H., 2nd, Hughes, T.K., Sutton, M.S., Nyquist, S.K., Bromley, J.D., Mondal, R., et al. (2025). High-dose intravenous BCG vaccination induces enhanced immune signaling in the airways. Sci Adv 11, eadq8229. 10.1126/sciadv.adq8229.

23. Maiello, P., DiFazio, R.M., Cadena, A.M., Rodgers, M.A., Lin, P.L., Scanga, C.A., and Flynn, J.L. (2018). Rhesus Macaques Are More Susceptible to Progressive Tuberculosis than Cynomolgus Macaques: a Quantitative Comparison. Infect Immun 86. 10.1128/IAI.00505-17.

24. Darrah, P.A., Zeppa, J.J., Wang, C., Irvine, E.B., Bucsan, A.N., Rodgers, M.A., Pokkali, S., Hackney, J.A., Kamath, M., White, A.G., et al. (2023). Airway T cells are a correlate of i.v. Bacille Calmette-Guerin-mediated protection against tuberculosis in rhesus macaques. Cell Host Microbe 31, 962–977 e968. 10.1016/j.chom.2023.05.006.

25. Ganchua, S.K.C., Cadena, A.M., Maiello, P., Gideon, H.P., Myers, A.J., Junecko, B.F., Klein, E.C., Lin, P.L., Mattila, J.T., and Flynn, J.L. (2018). Lymph nodes are sites of prolonged bacterial persistence during Mycobacterium tuberculosis infection in macaques. PLOS Pathogens 14, e1007337. 10.1371/journal.ppat.1007337.

26. Ganchua, S.K.C., White, A.G., Klein, E.C., and Flynn, J.L. (2020). Lymph nodes—The neglected battlefield in tuberculosis. PLOS Pathogens 16, e1008632. 10.1371/journal.ppat.1008632.

27. Jayalakshmi, P., Malik, A.K., and Soo-Hoo, H.S. (1994). Histopathology of lymph nodal tuberculosis--university hospital experience. Malays J Pathol 16, 43–47.

28. Mattila, J.T., Maiello, P., Sun, T., Via, L.E., and Flynn, J.L. (2015). Granzyme B-expressing neutrophils correlate with bacterial load in granulomas from Mycobacterium tuberculosis-infected cynomolgus macaques. Cell Microbiol 17, 1085–1097. 10.1111/cmi.12428.

29. McCaffrey, E.F., Delmastro, A.C., Fitzhugh, I., Ranek, J.S., Douglas, S., Peters, J.M., Fullaway, C.C., Bosse, M., Liu, C.C., Gillen, C., et al. (2026). The immunometabolic topography of cellular organization and bacterial control in tuberculosis granulomas. Nat Immunol 27, 867–880. 10.1038/s41590-026-02431-8.

30. Lin, P.L., Ford, C.B., Coleman, M.T., Myers, A.J., Gawande, R., Ioerger, T., Sacchettini, J., Fortune, S.M., and Flynn, J.L. (2014). Sterilization of granulomas is common in active and latent tuberculosis despite within-host variability in bacterial killing. Nat Med 20, 75–79. 10.1038/nm.3412.

31. Ordway, D.J., Shang, S., Henao-Tamayo, M., Obregon-Henao, A., Nold, L., Caraway, M., Shanley, C.A., Basaraba, R.J., Duncan, C.G., and Orme, I.M. (2011). Mycobacterium bovis BCG-mediated protection against W-Beijing strains of Mycobacterium tuberculosis is diminished concomitant with the emergence of regulatory T cells. Clin Vaccine Immunol 18, 1527–1535. 10.1128/CVI.05127-11.

32. Peters, N.C., Pagan, A.J., Lawyer, P.G., Hand, T.W., Henrique Roma, E., Stamper, L.W., Romano, A., and Sacks, D.L. (2014). Chronic parasitic infection maintains high frequencies of short-lived Ly6C+CD4+ effector T cells that are required for protection against re-infection. PLoS Pathog 10, e1004538. 10.1371/journal.ppat.1004538.

33. Roy Chowdhury, R., Vallania, F., Yang, Q., Lopez Angel, C.J., Darboe, F., Penn-Nicholson, A., Rozot, V., Nemes, E., Malherbe, S.T., Ronacher, K., et al. (2018). A multi-cohort study of the immune factors associated with M. tuberculosis infection outcomes. Nature 560, 644–648. 10.1038/s41586-018-0439-x.

34. Chen, C.Y., Huang, D., Wang, R.C., Shen, L., Zeng, G., Yao, S., Shen, Y., Halliday, L., Fortman, J., Mcallister, M., et al. (2009). A Critical Role for CD8 T Cells in a Nonhuman Primate Model of Tuberculosis. PLoS Pathogens 5, e1000392. 10.1371/journal.ppat.1000392.

35. Smith, A.A., Su, H., Wallach, J., Liu, Y., Maiello, P., Borish, H.J., Winchell, C., Simonson, A.W., Lin, P.L., Rodgers, M., et al. (2025). A BCG kill switch strain protects against Mycobacterium tuberculosis in mice and non-human primates with improved safety and immunogenicity. Nat Microbiol 10, 468–481. 10.1038/s41564-024-01895-4.

36. Hughes, T.K., Wadsworth, M.H., 2nd, Gierahn, T.M., Do, T., Weiss, D., Andrade, P.R., Ma, F., de Andrade Silva, B.J., Shao, S., Tsoi, L.C., et al. (2020). Second-Strand Synthesis-Based Massively Parallel scRNA-Seq Reveals Cellular States and Molecular Features of Human Inflammatory Skin Pathologies. Immunity 53, 878–894 e877. 10.1016/j.immuni.2020.09.015.

37. Gierahn, T.M., Wadsworth, M.H., 2nd, Hughes, T.K., Bryson, B.D., Butler, A., Satija, R., Fortune, S., Love, J.C., and Shalek, A.K. (2017). Seq-Well: portable, low-cost RNA sequencing of single cells at high throughput. Nat Methods 14, 395–398. 10.1038/nmeth.4179.

38. Macosko, E.Z., Basu, A., Satija, R., Nemesh, J., Shekhar, K., Goldman, M., Tirosh, I., Bialas, A.R., Kamitaki, N., Martersteck, E.M., et al. (2015). Highly Parallel Genome-wide Expression Profiling of Individual Cells Using Nanoliter Droplets. Cell 161, 1202–1214. 10.1016/j.cell.2015.05.002.

39. Li, B., Gould, J., Yang, Y., Sarkizova, S., Tabaka, M., Ashenberg, O., Rosen, Y., Slyper, M., Kowalczyk, M.S., Villani, A.C., et al. (2020). Cumulus provides cloud-based data analysis for large-scale single-cell and single-nucleus RNA-seq. Nat Methods 17, 793–798. 10.1038/s41592-020-0905-x.

40. Dobin, A., Davis, C.A., Schlesinger, F., Drenkow, J., Zaleski, C., Jha, S., Batut, P., Chaisson, M., and Gingeras, T.R. (2013). STAR: ultrafast universal RNA-seq aligner. Bioinformatics 29, 15–21. 10.1093/bioinformatics/bts635.

41. Andreatta, M., Corria-Osorio, J., Muller, S., Cubas, R., Coukos, G., and Carmona, S.J. (2021). Interpretation of T cell states from single-cell transcriptomics data using reference atlases. Nat Commun 12, 2965. 10.1038/s41467-021-23324-4.

42. Capuano, S.V., Croix, D.A., Pawar, S., Zinovik, A., Myers, A., Lin, P.L., Bissel, S., Fuhrman, C., Klein, E., and Flynn, J.L. (2003). Experimental *Mycobacterium tuberculosis* Infection of Cynomolgus Macaques Closely Resembles the Various Manifestations of Human *M. tuberculosis* Infection. Infection and Immunity 71, 5831–5844. 10.1128/iai.71.10.5831-5844.2003.

43. Lin, P.L., Rodgers, M., Smith, L.K., Bigbee, M., Myers, A., Bigbee, C., Chiosea, I., Capuano, S.V., Fuhrman, C., Klein, E., and Flynn, J.L. (2009). Quantitative Comparison of Active and Latent Tuberculosis in the Cynomolgus Macaque Model. Infection and Immunity 77, 4631–4642. 10.1128/iai.00592-09.

44. Sarnyai, Z., Nagy, K., Patay, G., Molnár, M., Rosenqvist, G., Tóth, M., Takano, A., Gulyás, B., Major, P., Halldin, C., and Varrone, A. (2019). Performance Evaluation of a High-Resolution Nonhuman Primate PET/CT System. Journal of Nuclear Medicine 60, 1818–1824. 10.2967/jnumed.117.206243.

45. Hartman, A.L., Nambulli, S., McMillen, C.M., White, A.G., Tilston-Lunel, N.L., Albe, J.R., Cottle, E., Dunn, M.D., Frye, L.J., Gilliland, T.H., et al. (2020). SARS-CoV-2 infection of African green monkeys results in mild respiratory disease discernible by PET/CT imaging and shedding of infectious virus from both respiratory and gastrointestinal tracts. PLoS Pathog 16, e1008903. 10.1371/journal.ppat.1008903.

46. Lin, P.L., Maiello, P., Gideon, H.P., Coleman, M.T., Cadena, A.M., Rodgers, M.A., Gregg, R., O’Malley, M., Tomko, J., Fillmore, D., et al. (2016). PET CT Identifies Reactivation Risk in Cynomolgus Macaques with Latent M. tuberculosis. PLoS Pathog 12, e1005739. 10.1371/journal.ppat.1005739.

47. Darrah, P., Zeppa, J., Maiello, P., Hackney, J., Wadsworth, M., Hughes, T., Pokkali, S., Swanson, P., Grant, N., Rodgers, M., et al. (2020). Prevention of tuberculosis in macaques after intravenous BCG immunization. Nature 577, 95–102. 10.1038/s41586-019-1817-8.

