## Supplemental Figures S1-S11 for "Protection against disseminated disease following reinfection with *Mycobacterium tuberculosis* requires CD8+ lymphocytes"

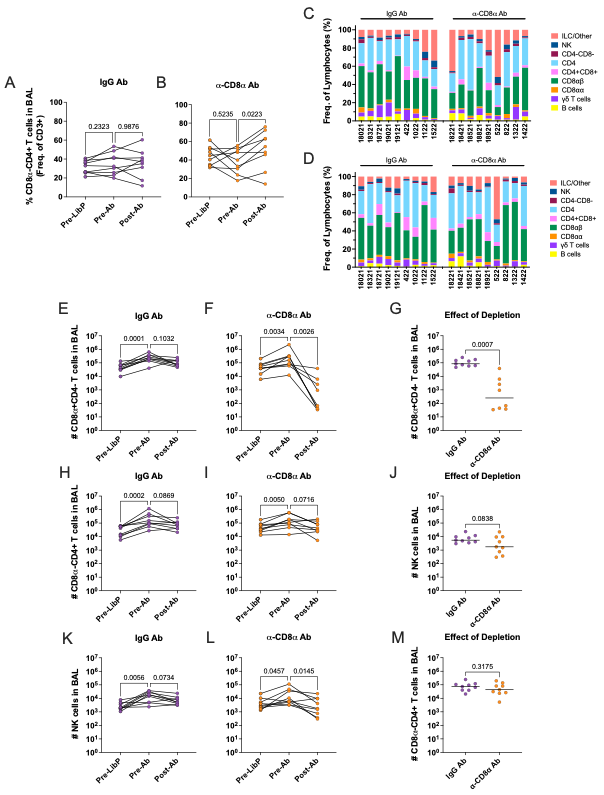
 **Supplemental Figure S1. Impact of primary infection and antibody depletion on airway lymphocyte composition.** (**A-B**) Frequency of CD4+ T cells over time shown as a proportion of CD3+ lymphocytes for animals receiving IgG (A) or α-CD8α antibody (B). (**C-D**) Relative abundance of lymphocyte subsets (**C**) prior to primary (i.e. LibP) infection and (**D**) prior to starting depletion infusions. Each bar represents a BAL from one animal. (**E-M**) Quantification of select lymphocyte subsets in the airway throughout the primary infection. (**E-G**) CD8a+ T cells. (**H-J**) NK cells. (**K-M**) CD4+ T cells. Serial quantification of cell types for animals receiving (**E,H,K**) IgG isotype or (**F,I,L**) α-CD8α depleting antibodies. (**G,J,M**) Comparison of IgG and α-CD8α groups following depletion. Each symbol represents an animal; thin lines connect animals across timepoints, thick horizontal lines represent group median. For panels A-B,E-F,H-I,K-L, p-values shown from repeated measures one-way ANOVA with Dunnett's multiple comparisons test. For panels G,J,M, p-values shown from Welch’s t test.

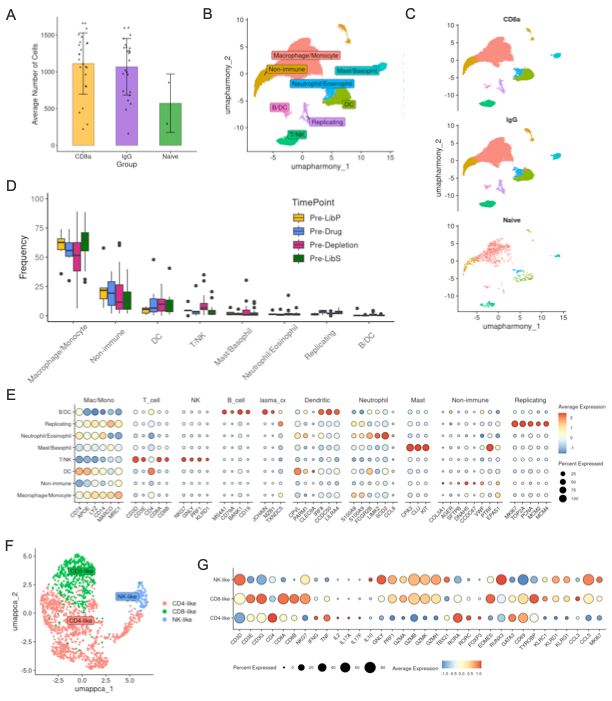

**Supplemental Figure S2. scRNAseq identification and analysis T cell populations in airways.** (**A**) Number of cells recovered per sample during sequencing. Each dot represents a BAL sample, colored bars represent mean per group and error bars show standard deviation. (**B**) UMAP of coarse level cell type annotation. (**C**) Coarse level populations shown across depletion group. (**D**) Relative frequency of each coarse level population shown across each of four timepoints. (**E**) Dot plot of common coarse level genes for annotated cell types. (**F**) UMAP of T/NK cell subset annotation. (**G**) Dot plot of common T/NK genes for subsets identified via subclustering.

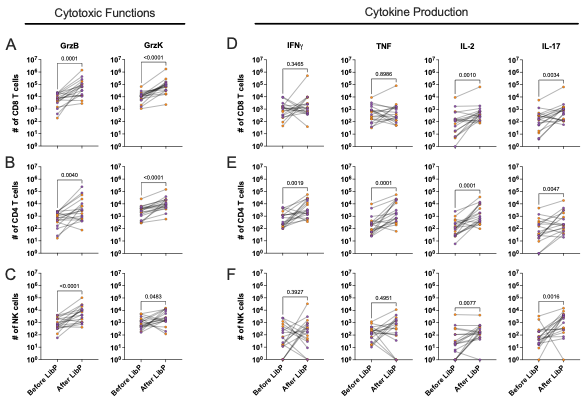

**Supplemental Figure S3. Lymphocyte function in airway following previous infection with Mtb.** (**A-F**) Number of (**A,D**) CD8αβ T cells, (**B,E**) CD4 T cells, and (**C,F**) NK cells producing (**A-C**) cytotoxic effector molecules and (**D-F**) cytokines in BAL. Each symbol represents an animal; lines connect animals across timepoints. P-values shown from Wilcoxon signed-rank non-parametric test.

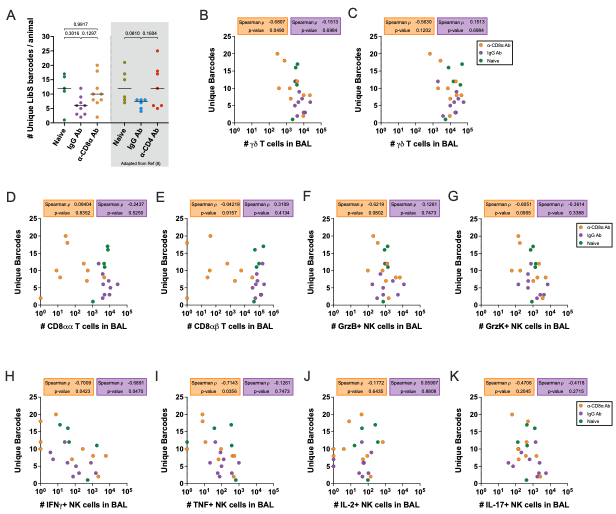

**Supplemental Figure S4. NK cell control of infection establishment.** (**A**) Number of unique barcodes recovered from CFU+ tissues at necropsy. In CD8α depletion study, all groups were compared against each other. In CD4 study, all groups were compared to the non-depleted IgG reinfection control. P values shown are for Welch’s ANOVA with Dunnett’s T3 multiple comparisons test. Each symbol represents an animal; line represents group median. (**B-E**) Relationship between the number of (B) NK cells, (C) γδ T cells, (D) CD8αα T cells and (E) CD8αβ T cells in the airway prior to LibS infection and the number of unique barcodes recovered at necropsy. (**F-K**) Relationship between the number of (F) granzyme B, (G) granzyme K, (H) IFN-γ, (I) TNF, (J) IL-2, and (K) IL-17 producing NK cells in the airway prior to infection and the number of unique barcodes recovered at necropsy. For panels B-K, Spearman rho (ρ) and p value reported for each correlation in α-CD8α Ab (orange) and IgG Ab (purple) groups. No statistical analysis was performed on the naïve (green) dataset due to small sample size.

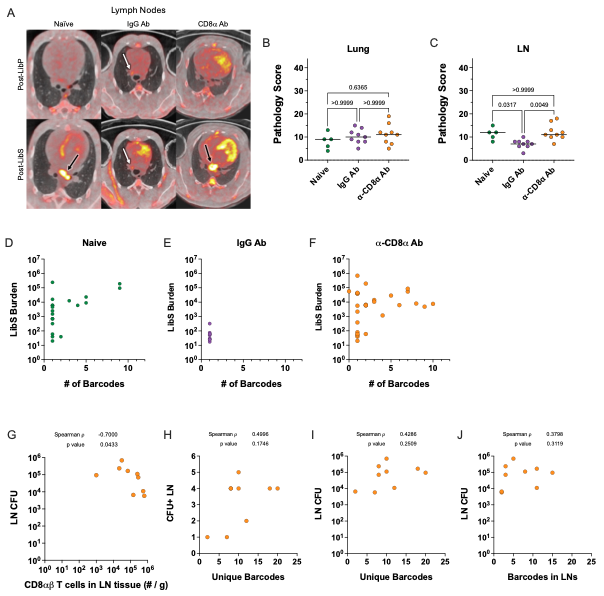

**Supplemental Figure S5. Analysis of early dissemination events across models of protection.** (**A**) Representative slices from PET CT scans demonstrating annotation of involved lymph nodes, temporal tracking with serial scans and differentiating new and old sites of infection. White arrow indicates an “old” lymph node involved in the primary LibP infection. Black arrows indicate “new” lymph nodes that became inflamed after the secondary LibS infection. (**B,C**) Pathology scores for (**B**) lung and (**C**) LN compartments. P values are for Kruskal-Wallis non-parametric test with multiple comparisons. Each symbol represents an animal; line represents group median. (**D-F**) Relationship between the number of barcodes identified in a LN and the bacterial burden within that tissue for (**D**) naïve, (**E**) IgG and (**F**) CD8α depleted animals. Each symbol represents a lymph node. (**G**) Relationship between the number of CD8αβ T cells per gram of LN tissue and the total LibS bacterial burden in LN. (**H**) Relationship between the overall number of unique barcodes identified in an animal and the number of CFU+ LNs. (**I**) Relationship between the overall number of unique barcodes identified in an animal and the total LibS bacterial burden in LN. (**J**) Relationship between the number of barcodes identified in LN tissues and the total LibS bacterial burden in LN. For panels D-G, Spearman rho (ρ) and p value reported for each correlation.

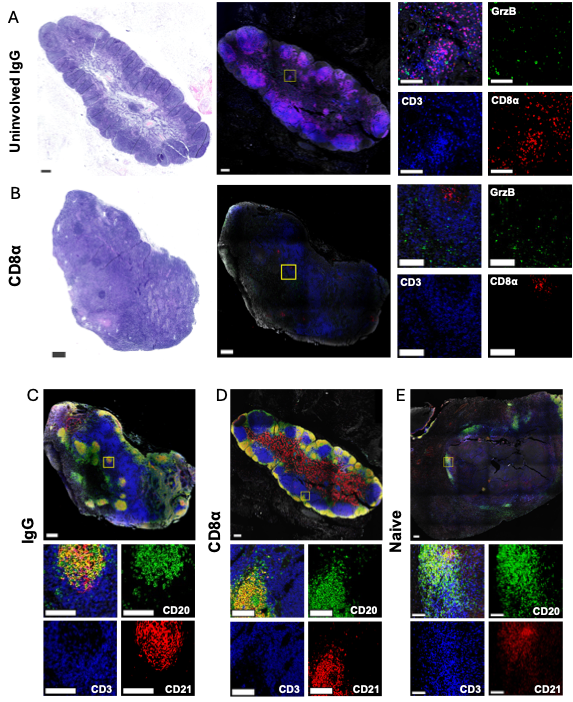

**Supplemental Figure S6. Uninvolved LN staining and identification of follicular DCs.** (**A,B**). H&E staining and DAPI (*grey*), CD8α (*red*), Granzyme B (*green*), and CD3 (*blue*) IFA for (**A**) an individual uninvolved peripheral LN from an IgG animal and (**B**) a cranial hilar LN from a depleted animal. Several CD8α+ clusters are notated with arrows. (**C-E**). DAPI (*grey*), CD21 (*red*), CD20 (*green*), and CD3 (*blue*) IFA on serial sections of (**C**) the same uninvolved IgG LN as in panel A, (**D**) the same depleted LN as in panel B, and (**E**) a naïve LN. For all large images, scale bars represent 300μM. For all inset images, scale bars represent 100μM.

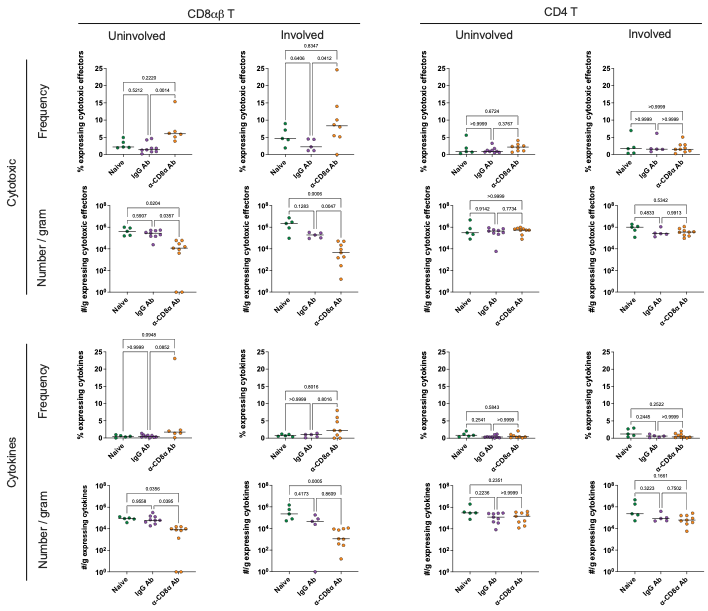

**Supplemental Figure S7. Function of CD8 and CD4 T cells in LNs.** Boolean analysis of intracellular cytokine staining (ICS) on LN tissues. Conventional CD8 (columns 1 and 2) and CD4 (columns 3 and 4) were assessed for production of cytotoxic effector molecules (rows 1 and 2) and cytokines (rows 3 and 4). Cytotoxic effectors evaluated included granulysin, perforin, and granzymes B, K and M. Cytokines evaluated included IFN-γ, TNF, IL-2 and IL-17. Frequency (rows 1 and 3) and number per gram of tissue (rows 2 and 4) of positive cells are shown for both uninvolved (columns 1 and 3) and involved (columns 2 and 4) tissues. For panels showing frequency, p values shown are for Kruskal-Wallis non-parametric test with Dunn’s multiple comparisons. For panels showing number of cells per gram, p values shown are from Welch’s ANOVA with Dunnett’s T3 multiple comparisons tests for pairwise comparisons of all three treatment groups. Samples below threshold for event counts were excluded from frequency analysis (see Methods).

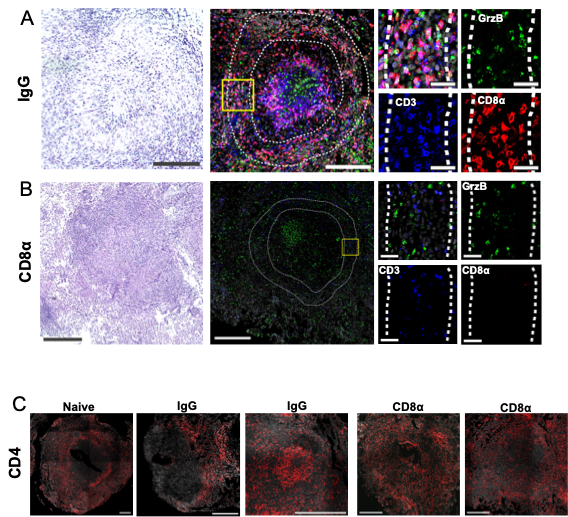

**Supplemental Figure S8. Additional lung granuloma IFA and quantification ROIs.** (**A-B**) Additional FFPE lung granulomas from (**A**) an IgG animal and (**B**) a depleted animal were stained with hematoxylin and eosin (H&E, *left* column) to characterize disease and tissue morphology followed by IFA staining (*middle*, *right* columns) with DAPI (*grey*), CD8α (*red*), Granzyme B (*green*) and CD3 (*blue*) to assess lymphocytes. (**C**) The same lung granulomas from panels A,B and Fig. 5A-C were re-stained in a cyclic staining protocol to identify CD4+ T cells (*red*). In panel A, the scale bars represent 150μM for large images and 30μM for insets. In panel B, scale bars represent 300μM for large images and 30 μM for insets. In panel C, all scale bars represent 300 μM.

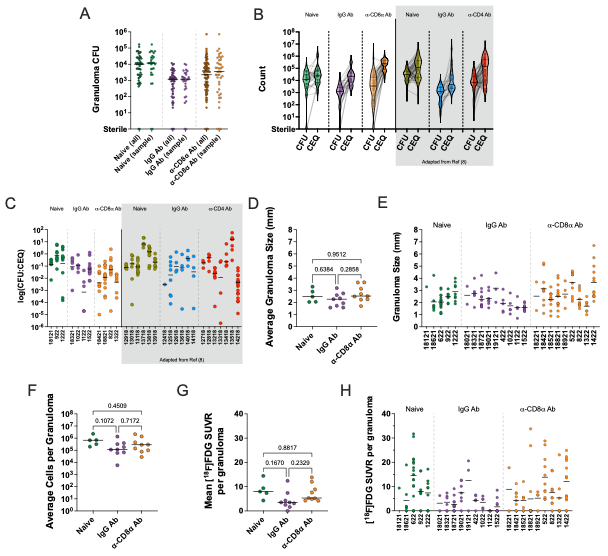

**Supplemental Figure S9. Granuloma anaylsis.** (**A**) Comparison of all granulomas from each treatment group to the subset used for CEQ analysis to ensure a representative sampling. Each symbol represents a granuloma, line represents group median. (**B**) Relationship between CFU and CEQ within individual lesions. Each symbol represents a granuloma, thin lines connect animals across time points and thick line in violin plot represents group median. (**C**) Killing capacity in individual lesions granulomas, shown by the ratio between CFU and CEQ. (**D,E**) Mean granuloma size, measured by PET CT scans. (**F**) The mean number of cells per granuloma, quantified by hemocytometer after homogenization. (**G,H**) Mean inflammation of granulomas quantified by PET signal. For panels, C,E,H, each symbol represents a granuloma, line represents animal mean. For panels, D,F,G, each symbol represents an animal, line represents group median. P values shown are for Welch’s ANOVA with Dunnett’s T3 multiple comparisons test.

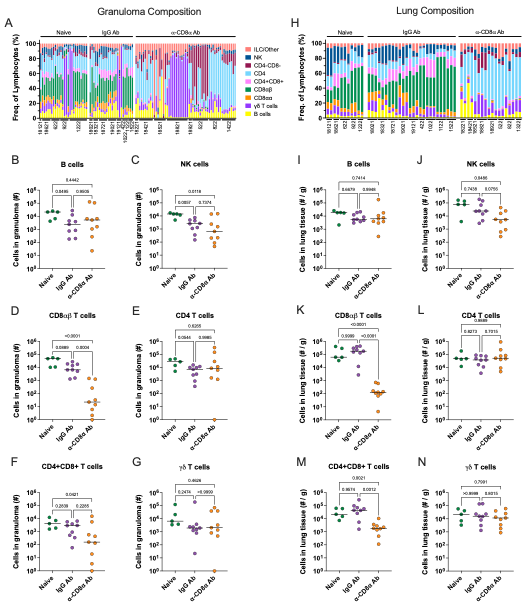

**Supplemental Figure S10. Lymphocyte composition of lung granulomas and tissue.** (**A**) Relative abundance of lymphocyte subsets in granulomas. Each bar represents a granuloma. (**B-G**) Number of (**B**) B cells, (**C**) NK cells, (**D**) CD8αβ T cells, (**E**) CD4 T cells, (**F**) CD4+CD8+ double positive T cells and (**G**) γδ T cells within granulomas. (**H**) Relative abundance of lymphocyte subsets in lung tissue. Each bar represents a randomly sampled lobe. (**I-N**) Number of (**I**) B cells, (**J**) NK cells, (**K**) CD8αβ T cells, (**L**) CD4 T cells, (**M**) CD4+CD8+ double positive T cells and (**N**) γδ T cells per gram of tissue. For panels B-G,I-N, p values shown are for Welch’s ANOVA with Dunnett’s T3 multiple comparisons test. Each symbol represents an animal; line represents group median.

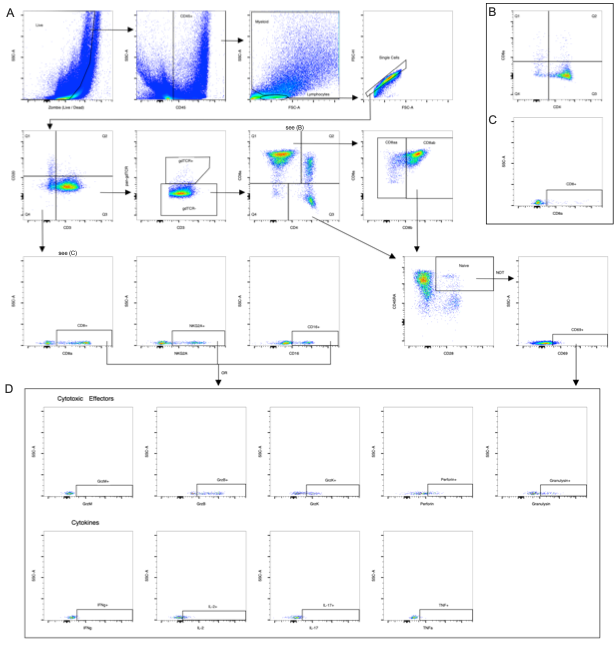

**Supplemental Figure S11. Gating strategy for flow cytometric analysis on representative sample.** (**A**) Identification of cell phenotype. (**B-C**) Representative flow plots showing depletion of CD8α+ T cells (B) and NK cells (C). (**D**) Identification of effector functions.
